# Colony pattern development of a synthetic bistable switch

**DOI:** 10.1101/2024.06.17.599191

**Authors:** Pan Chu, Jingwen Zhu, Zhixin Ma, Xiongfei Fu

## Abstract

Microbial colony development hinges upon a myriad of factors, including mechanical, biochemical, and environmental niches, which collectively shape spatial patterns governed by intricate gene regulatory networks. The inherent complexity of this phenomenon necessitates innovative approaches to comprehend and compare the mechanisms driving pattern formation. Here, we unveil the multistability of bacterial colony patterns orchestrated by a simple synthetic bistable switch. Utilizing quantitative imaging and spatially resolved transcriptome approaches, we explore the deterministic process of a ring-like colony pattern formation from a single cell. This process is primarily driven by bifurcation events programmed by the gene regulatory network and microenvironmental cues. Additionally, we observe a noise-induced process amplified by the founder effect, leading to patterns of symmetry-break during range expansion. The degrees of asymmetry are profoundly influenced by the initial conditions of single progenitor cells during the nascent stages of colony development. These findings underscore how the process of range expansion enables individual cells, exposed to a uniform growth-promoting environment, to exhibit inherent capabilities in generating emergent, self-organized behaviour.

## MAIN

Unraveling the formation of spatially structured living communities and the underlying mechanisms is a cornerstone in the fields of developmental biology, evolutionary biology, and ecology^1^. These patterns are evident in a wide array of contexts, from microbial biofilms and dense colonies^2,3^, to developing embryos^4^, differentiating tissues, and tumour progression ^5,6^. Mechanical and biochemical interactions, alongside processes like cell growth and migration, contribute to the establishment of spatial patterns^7–9^. Gene regulatory networks (GRNs), influenced by mechanical, biochemical, and environmental factors, govern this intricate orchestration. Additionally, stochasticity introduces a layer of randomness, contributing to variability in the system’s behaviour and adding to the observed diversity in biological outcomes^10–13^. As a result, complex organization emerges across various spatial and temporal scales.^2,14^ Understanding the underling mechanisms requires a thorough examination of coordinated interactions across multiple scales, encompassing molecular, cellular, and population levels. This multiscale complexity makes it challenging to elucidate the core principles of spatial patterning, necessitating alternative approaches capable of interrogating and comparing different pattern formation mechanisms. The rise of synthetic biology has provided a platform to construct synthetic systems that can explore these core patterning principles^15–18^.

In this study, we demonstrate the pattern diversity of bacterial colony orchestrated by a simple bistable genetic circuit in a system undergoing range expansion (Fig. 1a). We investigate the development of dense bacterial colonies from single cells, revealing that colony development follows a deterministic process influenced by GRNs and the microenvironment. In addition, a noise-induced process, amplified by the founder effect^19^, causes symmetry-breaking during expansion. These insights highlight how range expansion enables individual cells in a uniformly growth-promoting environment to exhibit emergent, self-organized behaviour.

**Fig. 1:**
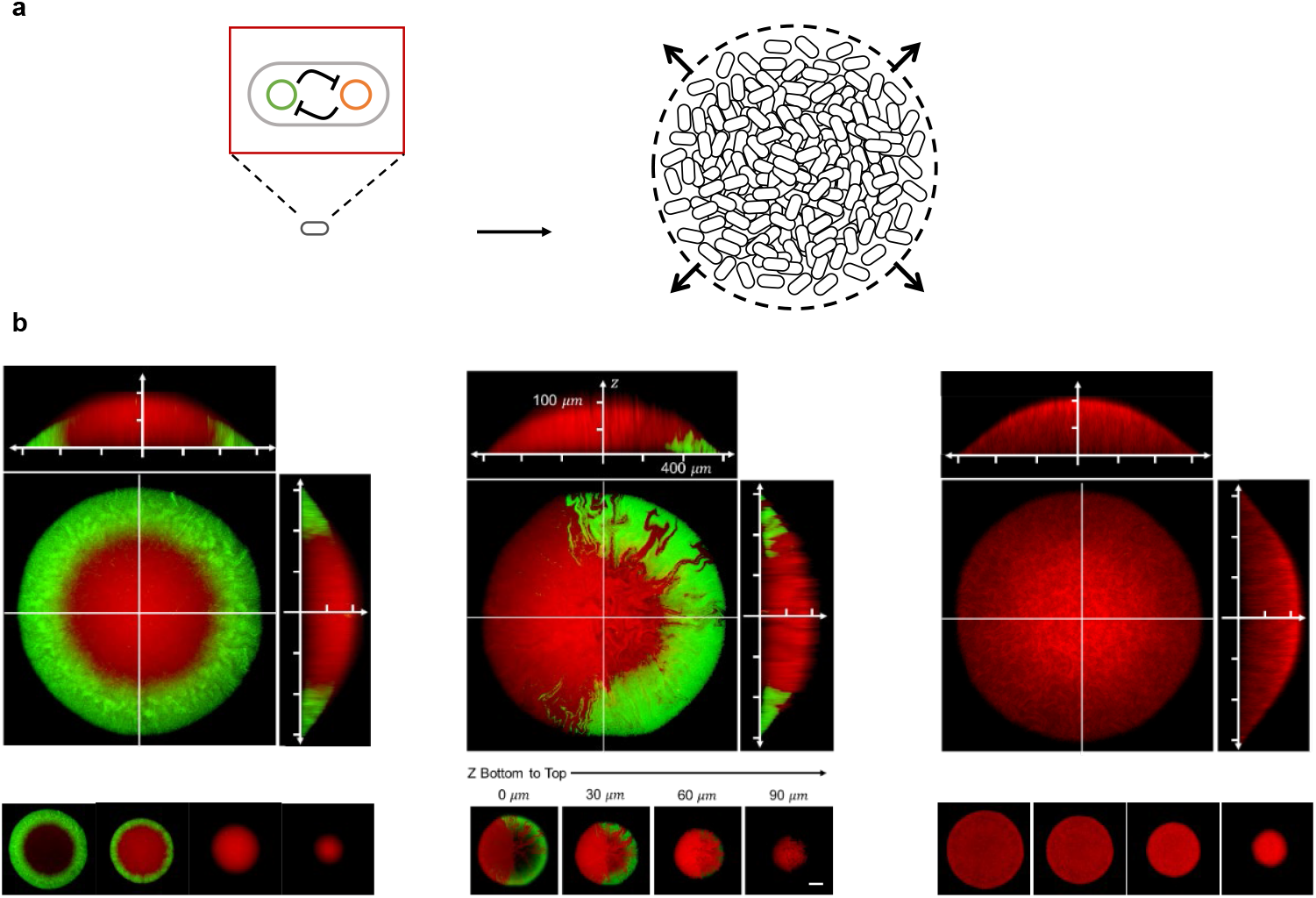
Spatial heterogeneity within a dense bacterial colony originating from a single cell during range expansion. (a) The non-motile *E. coli* strain, with a mutually repressive genetic circuit (depicted in Extended Data Fig. 1a), initiates growth from a single cell on 1.5% agar to form a dense colony. The mutual repression enables the bacterial cells to exhibit bistability, characterized by two distinct states: a red state with high expression of RFP and a green state with high expression of GFP. As cells in the colony proliferate and divide, those in the interior exert outward pressure, driving the expansion of the colony’s range. (b) Three representative colonies formed after 13 hours incubation (see Methods and Extended Data Fig. 1). Maximum intensity projection images are provided, along with the cross-sectional views of each colony, indicated at the top and right sides of the respective images. A gallery displaying each pattern is presented at the bottom of the images, with a scale bar indicating 200 µm.

### Diverse patterns emerge from a single cell

We grew the bacterial colony of non-motile engineered *E. coli* from a single cell on a solid agar plate (1.5% w/v) (Methods and Extended Data Fig. 1c and 1d). The bacterial cells are engineered by introduction of a well-characterized synthetic bistable switch^20–22^, which consists of two distinct phenotypic states denoted as the green state and red state corresponding to their respective fluorescence reporters (Extended Data Fig. 1a). Due to the mutually repressive topology of the genetic regulation, the engineered cells can maintain their respective states within the exponentially growing batch culture, unless the engineered system is toggled by two distinct inducers (Methods and Extended Data Fig. 1b). However, following an incubation period of 13 hours, we observed spontaneous development of the dense colony into a population comprising mixed phenotypic states. The spatial structures of the bacterial colony were categorized into three typical types: a ring-like pattern, a homogeneous one, and a sector-like pattern characterized by spatial segregation in phenotypes (Fig. 1b and Fig. S6).

We further employed a two-photon microscope to visualize the detailed spatial features within the densely populated colony associated with each type of pattern. The first type, a prominent ring-like pattern, comprised cells highly expressing GFP in the outer region of the colony, while cells highly expressing RFP were primarily located in the interior and upper regions (Fig. 1b left panel). The z-stack features from the bottom to the top of the colony, along with the cross-sectional perspective, indicated that the ring-like zone had approximately 200 µm in width and 50 µm in height. In addition, through flow cytometric analysis of cells picked from different regions of the colony, we confirmed that green state cells were located at the periphery of the colony, while red state cells were primarily on the top, along with some intermediate states (Extended Data Fig. 2a and 2g).

The second type of pattern was characterized by consistent red state cells throughout the entire colony. Significant variations in fluorescence intensity through the height of the colony were observed, with fluorescence intensity increasing with colony height (Fig. 1b, right panel, and Extended Data Fig. 2c, 2f and 2i).

The third type of pattern could be best described as a hybrid between the first and the second patterns. This configuration revealed a sectorial zonation, where the peripheral regions of the colony exhibited either the green or red state. The interior and upper regions of the colony consistently maintained the red state, irrespective of the state prevalent in the peripheral regions (Fig. 1b middle panel and Extended Data Fig. 2b, 2e and 2h).

The aforementioned observations suggested the spontaneous emergence of colony pattern diversity facilitated by a simple bistable switch. The underlying mechanism of pattern development, distinct from cell growth in a batch culture, intertwines with the complex dynamics of gene expression and cellular physiology. This prompts a fundamental question: How can such a relatively simple mutual repressive network direct individual cells into spatial patterns, thereby adopting distinct fates?

### Microenvironment heterogeneity reshapes the phenotype landscape

To better understand the developmental dynamics of various patterns originating from a single cell, we then tightly controlled the initial state of the cell. Interestingly, colonies initiated from green state cells exhibited exclusively the ring-like pattern (Fig. S6a). The emergence of red state cells was not observed in batch growth where cell growth gradually decreased during nutrient depletion (Extended Data Fig. 3).

Consequently, we postulated that the cell-fate landscape undergoes spatial and temporal dynamics throughout colony establishment (Fig. 2a). In a prior investigation, we revealed that the mutually repressive system experiences a growth-rate induced bifurcation, transitioning from bistability to monostability^22^. During colony development, the growth rate of individual cells at the local level, influenced by the microenvironmental factors^23–27^ such as local cell density^23^, nutrient availability^24–26^, oxygen access^28,29^, and mechanical effects^30^, exhibits both spatial and temporal variations within a densely populated colony. These variations play a crucial role in guiding subpopulations of cells within colonies toward distinct metabolic and physiological states. This dynamic progression of cell fates leads to the emergence of distinct states within the colony.

**Fig. 2:**
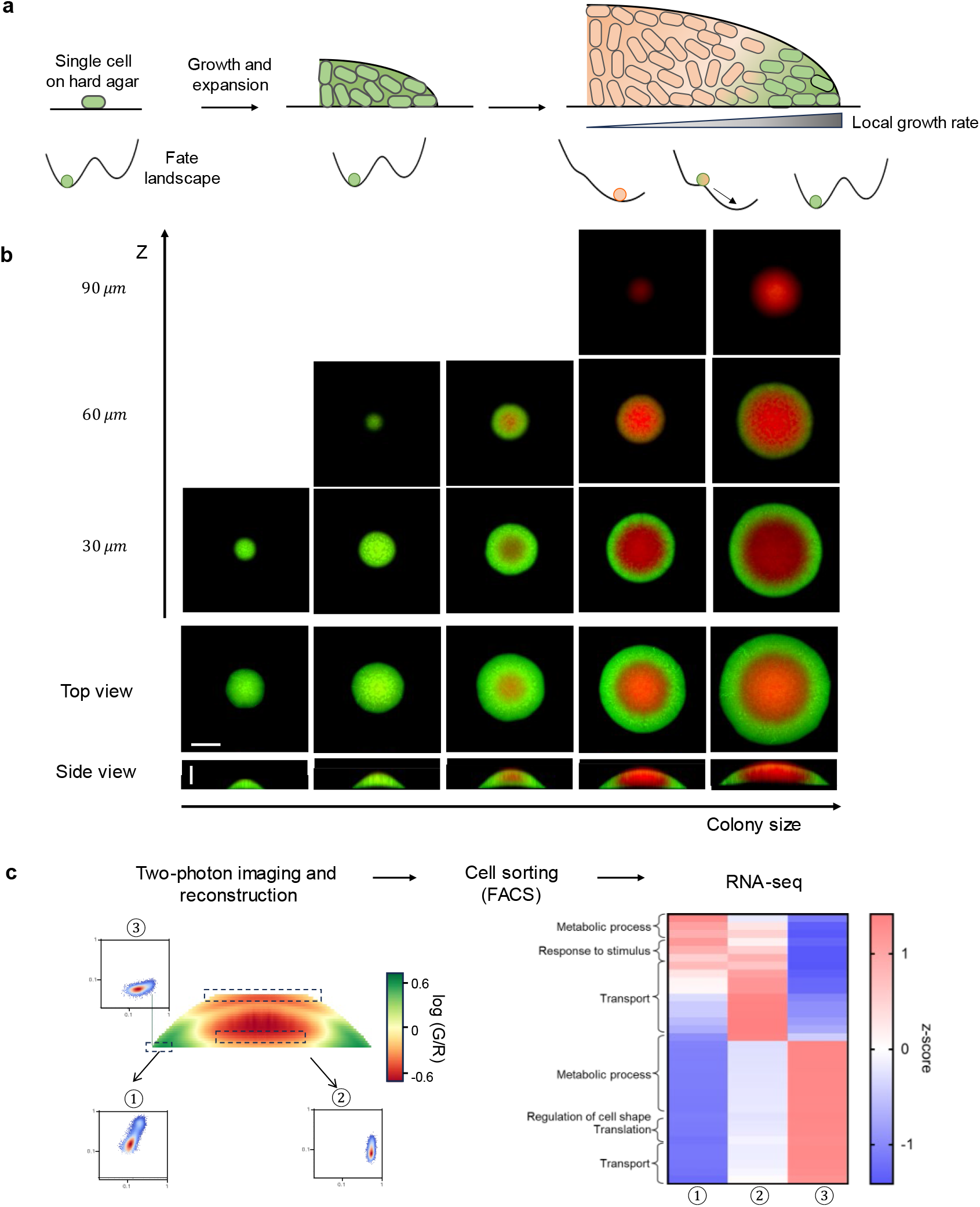
Microenvironment heterogeneity reshapes the phenotype landscape. (a) Spatial and temporal dynamics of cell fate landscapes throughout colony establishment, beginning from a single cell that rapidly divides to form a dense cluster. During the initial period after seeding, cells grow exponentially while the system remains bistable. As the colony expands, spatial heterogeneity emerges within the microenvironment, leading to spontaneous state transitions driven by variations in cell growth rates. This spatial heterogeneity and the resulting state transitions create a distinctive ring-like pattern within the colony. (b) Spatial distributions of colonies from 9 to 15 hours of growth post-seeding. The corresponding fluorescent distributions of cells in entire colonies of different sizes are provided in Extended Data Fig. 2. The horizontal scale bar represents 400 µm, while the vertical scale bar indicates 100 µm. (c) The framework of spatially resolved transcriptome analysis includes two-photon imaging and reconstruction, cell sorting according to fluorescence intensity distributions, and RNA-seq analysis. This analysis reveals a heterogeneous distribution of cell subpopulations with distinct metabolic functions. The functional landscape of the genes is presented (see details in Table S5).

To provide a comprehensive analysis of the phenotypic changes occurring within developing colonies, we tracked the dynamics of morphology and the distribution of cell states across colonies. From 9 to 16 hours after seeding, during the period referred to as the establishment phase^23^, the colony radius increased linearly with time at a constant radial speed of 120.1 µm/h, covering a range of radii from 200 to 1000 μm (Extended Data Fig. 4a). Consequently, we imaged colonies of various sizes to monitor the spatial-temporal dynamics of colony establishment (Fig. 2b). During the early stages, cells experience exponential growth, benefiting from the abundant availability of nutrients and oxygen. This robust growth is evident in the first and second columns of Fig. 2b, depicting colonies with radii less than 300 μm, where all cells remained in the green state. This observation aligns with the results of cytometry analysis of individual colonies (Extended Data Fig. 4e and 4f), confirming the presence of a uniform population in green state within the initial 10 hours following cell seeding. Subsequently, as the culture time progressed, a transient transition towards a heterogeneous population was observed. A portion of cells was observed to gradually transition to the red state as the colony continued to expand both radially and vertically (Fig. 2b and Extended Data Fig. 4e). The dynamics of the ring-like pattern development suggest that the spontaneous switch of cell phenotypes from the green state to the red state resulted from the variations in the colony microenvironment.

To further elucidate the effect of colony microenvironment on local physiological states, we generated a spatially resolved transcriptome for a dense colony (see Methods). We first reconstructed the spatial distribution of fluorescence intensity using z-stacks obtained from two-photon microscopy (Fig. 2c left panel). Subsequently, we sorted bacterial cells based on their spatial location in the colony using fluorescence-activated cell sorting (FACS, see Methods and Fig. S7). The transcriptomes confirmed that cells within distinct subpopulations exhibit variations in metabolic functions, distributed across different regions of the colony.^26,31^ Functional analysis of genes from different spatial regions reveals distinct patterns of gene expression within the colony. Specifically, genes associated with translation, amino acid biosynthesis, and nucleotide biosynthesis are upregulated in the colony interior and top layers. In contrast, genes involved in responding to environmental stimuli and across membrane transporters are predominantly upregulated in the colony periphery (Fig. 2c right panel and Table S5), indicating a heightened responsiveness to external signals and abundant nutrient access in the outer layers of the colony. Such spatial heterogeneity in the transcriptomes was observed to be independent of the specific growth medium (Fig. S8 and Table S6). Therefore, cells within the densely populated colony encountered highly heterogeneous microenvironments, which played a significant role in reshaping their phenotypes.

### Modulation of the ring width

To understand the determinants of colonization and the underlying mechanism of ring-like pattern formation, we deployed a continuum model^32,33^ of bacterial colonies that incorporate nutrient diffusion and uptake, growth of colony, and mechanical interactions between or within the colony and substrate (Supplementary Note 1). A detailed analysis of the model simulation indicates that the formation of the ring pattern is a deterministic process that is mainly governed by the bifurcation tipping point (the threshold at which state bifurcation occurs) within the mutual repressive network and the diffusion of substrate nutrient (Supplementary Note 1). Specifically, once the colony is established, it forms a steady traveling front expanding horizontally, accompanied by a steady traveling profile of nutrient distribution (Fig. S9). Further examination of nutrient distribution revealed that nutrient concentration falls below the corresponding threshold at both radial and vertical distances from the colony boundaries and agar surface (Fig. S9 and S10). In these regions, the system becomes monostable, triggering the state transition from the green state to the red state, thereby defining the boundary of green ring.

As the model suggested, the ring width of the green state cells remained constant during colony expansion (Fig. S11). By examining wide-field microscope images and calculating the ratio of GFP intensity to RFP intensity (GFP/RFP), we identified a distinct peak indicating the predominant GFP expression within the colony. This peak allowed us to define a green band width (Fig. 3a, 3b and Fig. S12), measured as the distance from the peak value to the colony edge. Remarkably, during the establishment phase of colony growth, characterized by a linear increase in colony radius over time, the green band width remained relatively constant (Fig. 3b, Extended Data Fig. 4a and 4d), indicating a robust stability in this characteristic feature of the ring-like pattern.

**Fig. 3:**
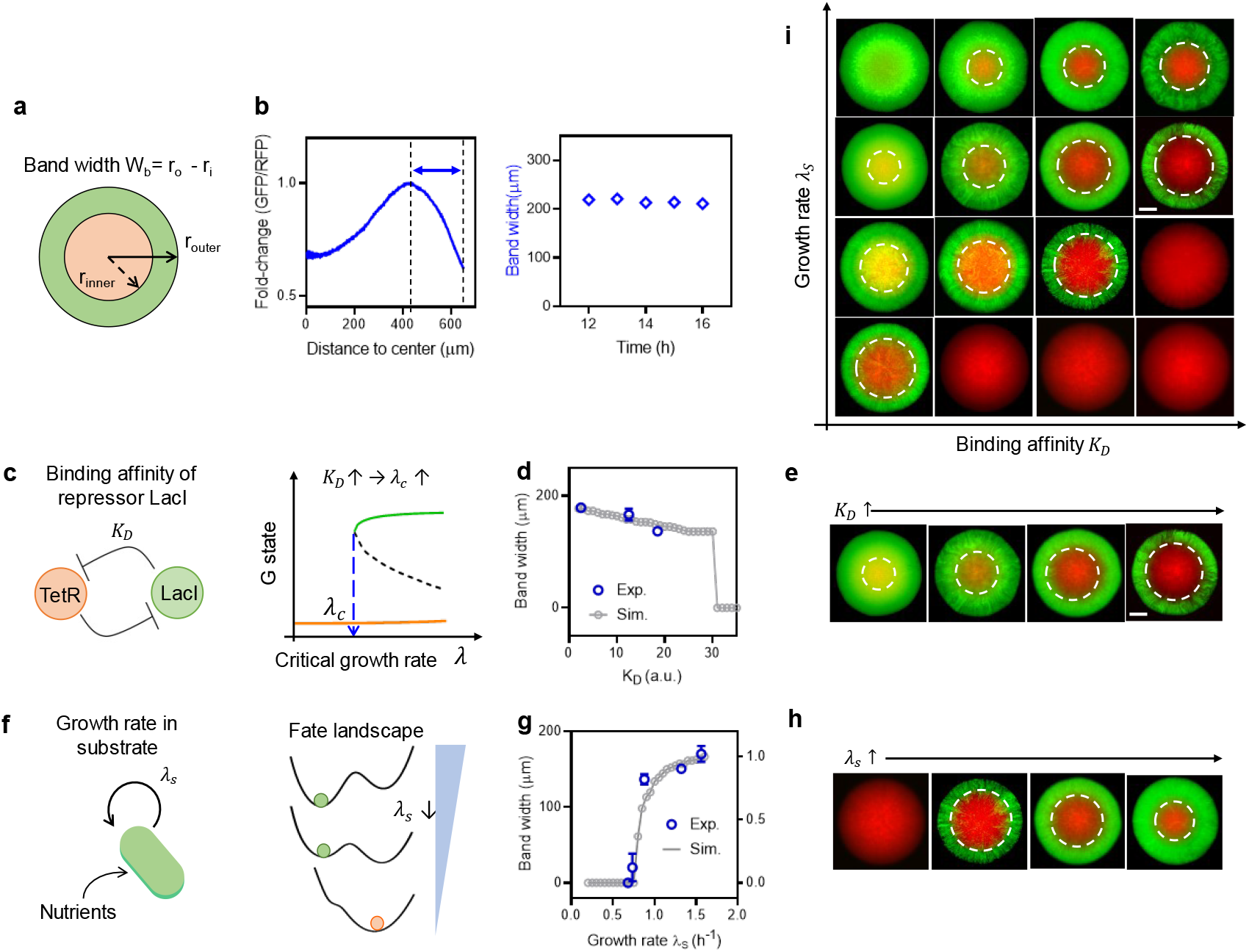
Modulation of the band width of the ring-like pattern. (a) The ring-like pattern is characterized by determining the band width where GFP expression predominates. (b) This is achieved by calculating the fold change, which is the ratio of GFP to RFP intensity, and identifying the maximum ratio, indicating predominant GFP expression (see Fig. S12). The band width was characterized as constant during the establishment phase, where the colony radius increases linearly (blue diamonds and Extended Data Fig. 4a). (c-e) Model prediction and experimental validation of the variation in band width by the critical growth rate (*λ_c_*) of different strains, where state transition occurs. (c) The critical growth rate is determined by the mutual repression network, which can be modulated by altering the binding affinity (*K_D_*) of the repressor (LacI) to its binding site, as detailed in Extended Data Fig. 5. (d) (e) Representative maximum intensity projection images are provided for different strains with diverse *K_D_* values grown on MOPS glucose medium agar plates. Scale bar indicates 200 µm. (f-h) Model prediction and experimental validation of the variation in band width by the maximal growth rate (*λ_s_*) determined by the quality of substrate nutrient. (g) (h) Representative images for the LO1 strain are shown under different conditions where the maximal growth rates range from 0.65 h^-1^ to 1.60 h^-1^ (see Table S7). (i) The phase diagram illustrates representative colony structures observed in the experiments by varying *K_D_* and *λ_s_*. The colonies presented in this figure were collected with a radius of approximately 600 µm.

Our model also predicted that the ring width was controlled by two key parameters: the critical transition growth rate *λ_c_* which is more specifically influenced by the binding affinity of the repressor LacI to its binding site *K_D_*, defining the bifurcation tipping point from monostability to bistability (Fig. 3c and 3d); and the maximum growth rate *λ_s_*, which defines the global nutrient quality of substrate media for the colony growth (Fig. 3f and 3g). Specifically, with an increasing *K_D_*, the thickness of the green state zone reduces both in vertical and radical distances, resulting in a smaller corresponding band width (Fig. 3d) On the other hand, the ring width decreased when the substrate nutrient quality *λ_s_* diminished (Fig. 3g). Further increase of *λ_c_* and decrease of *λ_s_* would lead the system experience a phase transition of complete loss of green state cells at the edge of the colony.

To validate the model prediction, we first manipulated the ring width by modifying the mutual repressive circuit, specifically altering the dissociation constant *K_D_* of LacI to its corresponding binding site (Fig. 3c). This modification exhibits varying critical transition growth rates *λ_c_* (Extended Data 5a, 5b and ref ^22^). Given that all these strains are genetically identical except for the lac operator, differing only by a few base pairs (Table S3), the growth rates of different strains and the spatiotemporal dynamics of colony establishment remained unchanged (Extended Data Fig. 5c and 5d). Assuming that the cell growth rate is proportional to the nutrient concentration, a higher *λ_c_* value indicates a larger nutrient concentration threshold below which the state transition occurs. Therefore, this resulted in a monotonic dependence of ring width on the *K_D_* (Fig. 3d).

Furthermore, we investigated the effect of global conditions for colony growth using different nutrient substrates, each providing a different maximum batch culture growth rate *λ_s_* (Table S7). As the nutrient quality becomes poorer, the ring width decreases (Fig. 3g and 3h). Interestingly, at certain lower *λ_s_* condition, where the mutual repression systems still exhibit a bistable feature (see ref^22^), cells within the colony were more prone to spontaneous transitions to the red state during the early stages of colony establishment, due to the heterogeneous distribution of nutrients resulting from consumption.

We systematically examined the features of colony patterns originating from single cells in the green state across all engineered mutants with different substrate media. The experimental results explicitly showed the bivariate dependency of the ring width and the pattern phase transition on *K_D_* and *λ_s_* (Fig. 3i and Fig. S13). Such dependence further indicated that the formation of ring pattern was a consequence of both internal regulations and external conditions.

### Noise-induced symmetry breaking in range expansion

Unlike the exclusively ring-like pattern observed in colonies initiated from green state cells, colonies starting from red state cells displayed all three types of patterns, as shown in Fig. 1b and Fig. S6b. To comprehend the pattern diversity initiated from the red state, particularly focusing on the diversity at the colony periphery, we monitored the early stage of pattern formation on an agar pad (see Methods and Extended Data Fig. 6). The microenvironment during early colony establishment can be considered as uniform. However, cells still experience fluctuations in gene expression that can induce cell fate transitions^12,34–38^. Due to the unbalanced feature of the mutual repression, the system tends to spontaneously transition into the green state (Extended Data Fig. 1b). Consequently, we observed instances where red cells switched to the green state, resulting in clusters of red and green cells during the early stages of colony development (Extended Data Fig. 6).

To further understand the cause and consequence of the phenotypic clusters, we expanded a previously reported agent-based model^23^ incorporating the biochemical network with stochasticity (see Supplementary Note 2). We monitored the cell colonization during the early time (t < 6 hours) forming only a thin layer of bacterial cells (Extended Data Fig. 7). Analysis of the simulation results led us to hypothesize that the observed symmetry-breaking pattern at the colony’s boundary, depicted in Fig. 1b, functions as a hallmark of range expansions featuring a moderate cell fate transition rate and range expansion rate (i.e., cell division rate). As shown in Fig. 4a, the distinct sectors, demonstrating different phenotypes within the ring-like zone of colonies, are believed to arise from the cell clusters formed within the initial hours after seeding (three types of clusters as illustrated in Fig. 4a right panel). The segregation is presumed to result from the founder effect within a narrow region of reproducing pioneers at the expanding frontier.^19^ In the later stages of colony establishment, with a higher density of founders, newly formed green state cells resulting from transitions from red ones become ensnared in the colony’s bulk, unable to further contribute to the colonization process on the agar plate.^19,39,40^ In another words, the stochastic model suggested that the types of patterns heavily depended on the state transition during the early colony development (Extended Data Fig. 7).

**Fig. 4:**
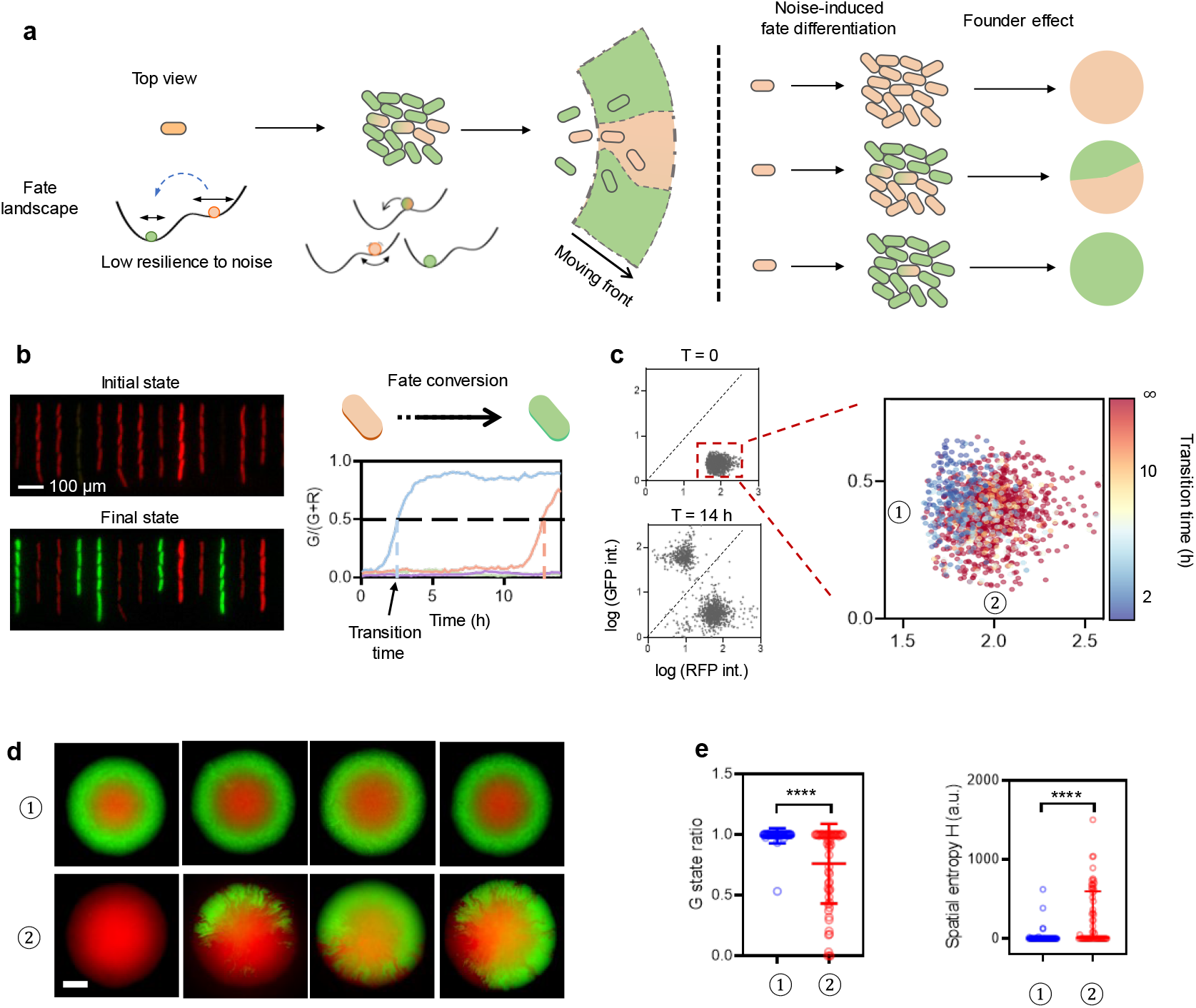
Noise-induced symmetry breaking in range expansion. (a) Visual representation of predicted scenarios where symmetry breaking occurs early in colony formation and is later amplified by the founder effect. (b) The mother machine microfluidic experiment was conducted under balanced conditions in RDM glucose medium (see Methods). Snapshots depict the initial state and the final state after 14 hours of culture. Scale bar indicates 100 µm. The first-passage time (right panel) indicates when the spontaneous state transition from the red state to the green state occurs, determined by the time point at which the fluorescent intensity ratio G/(G+R) for a given mother cell exceeds 0.5. (c) Density plot showing the distribution of fluorescence intensity among mother cells in the first recorded frame (T=0) and the final frame (T=14h). The first recorded frame plot provides insight into the initial state of the cells before the onset of the experiment. Two subpopulations, labeled as ① and ②, were selected, and the proportion of green cells was monitored over time for each subpopulation. In total, 1383 trajectories were recorded over a duration of 14 hours. The transition time from the red state to the green state for each mother cell is plotted in the right panel. Cells that did not switch their state within the 14-hour observation period are represented with an infinite transition time. Details on data processing, including cell segmentation and single cell tracking, are in the Methods. (d) Single-cell isolation by FACS (as illustrated in Extended Data Fig. 1d) enables the precise selection of individual cells for further analysis. By isolating cells from two different initial conditions (① and ② as indicated in (c)) and imaging the resulting colonies after 13 hours of incubation, different patterns were observed. Scale bar indicates 200 µm. (e) Features of bacterial colonies generated from different initial conditions. In the analysis of G state ratio and spatial entropy, bands with a width of 60 pixels, equivalent to 97.5 µm, positioned 300 to 360 pixels away from the colony center, were utilized (see Methods and Fig. S16). The lines in the plots indicate the median and interquartile ranges. To compare between groups, an unpaired, two-sided t-test with a 95% confidence interval was employed, revealing a significant difference (*\*\*\*\*P* < 2E-6). Additional colonies formed from single cells under different conditions are shown in Fig. S15 for further comparison.

We then investigated what determined the state transition probability by employing a microfluidic mother machine that enabled us to track individual red state cells under balanced growth, with an ample supply of the rich defined medium (see Methods). We found approximately 36% of red state mother cells transitioned to the green state (Fig. 4b and Fig. 4c left panel, see Methods), over a 14-hour cultivation period, corresponding to approximately 32 generations (doubling time ∼ 26 min). Despite all cells being in conditions where the RFP was highly expressed, they displayed significant heterogeneity in the trajectories of cell state transition from red state to green state (Fig. 4b right panel). For each trajectory, we defined the state transition time as the point at which the ratio of the GFP intensity (G/(G+R)) reached 0.5 (Fig. 4b right panel).

Interestingly, we observed that the state transition time was strongly related to the initial state of the cell being first tracked. Cells in subpopulations close to the green steady state (e.g., ① in Fig. 4c and Extended Data Fig. 8) exhibited much shorter transition times (Extended Data Fig. 8), indicating a higher probability of transitioning into the green state. In contrast, cells in subpopulations with higher RFP intensity (low GFP intensity, e.g., ② in Fig. 4c and Extended Data Fig. 8) preferentially remain in the red state (Extended Data Fig. 8). This observation suggests that even phenotypically homogeneous cell types exhibit a high degree of heterogeneity in the expression of individual genes, which may contribute to diverse potential for differentiation.

Given the knowledge about the diversity of the state transition time, we further experimentally tested the prediction by isolating single cells from a clonal population using FACS. Single cells in the red state from various subpopulations were sorted and then incubated on hard agars overnight to allow colonies to reach a radius of approximately 600 µm before being imaged using a fluorescent microscope (Fig. 4d). Cells sorted from the subpopulation proximate to the green state formed colonies with almost homogeneous GFP rings (① in Fig. 4d and Fig. S14), resembling colonies from the green state single cells. Divergent spatial configurations emerge from single cells sorted based on higher RFP expression levels (② in Fig. 4d and Fig. S14), exhibiting increased spatial entropy (Fig. 4d right panel and Methods). We also examined the effects of diverse state transition propensities and expansion speeds (Fig. S15), by various combinations of growth conditions and genetic circuits. This analysis illustrates the significance of both spontaneous state transition probability and range expansion velocity in the formation of pattern symmetry breaking.

## Discussion

Diversity in patterns is widely observed in nature, yet the underlying mechanisms remain poorly understood.^41,42^ The multiscale complexity of this process makes elucidating the core principles of spatial patterning challenging in living systems^43^. In this study, we discovered that a simple synthetic bistable switch could generate a variety of colony patterns originating from a single cell during the range expansion process. Our experimental and modelling results delineated the conditions and mechanisms under which deterministic bifurcation and noise-induced tipping contribute to the comprehensive formation of spatial patterns and multistability (Fig. 5). Our synthetic approach provides a promising example for interrogating and comparing different mechanisms of pattern formation.

**Fig. 5:**
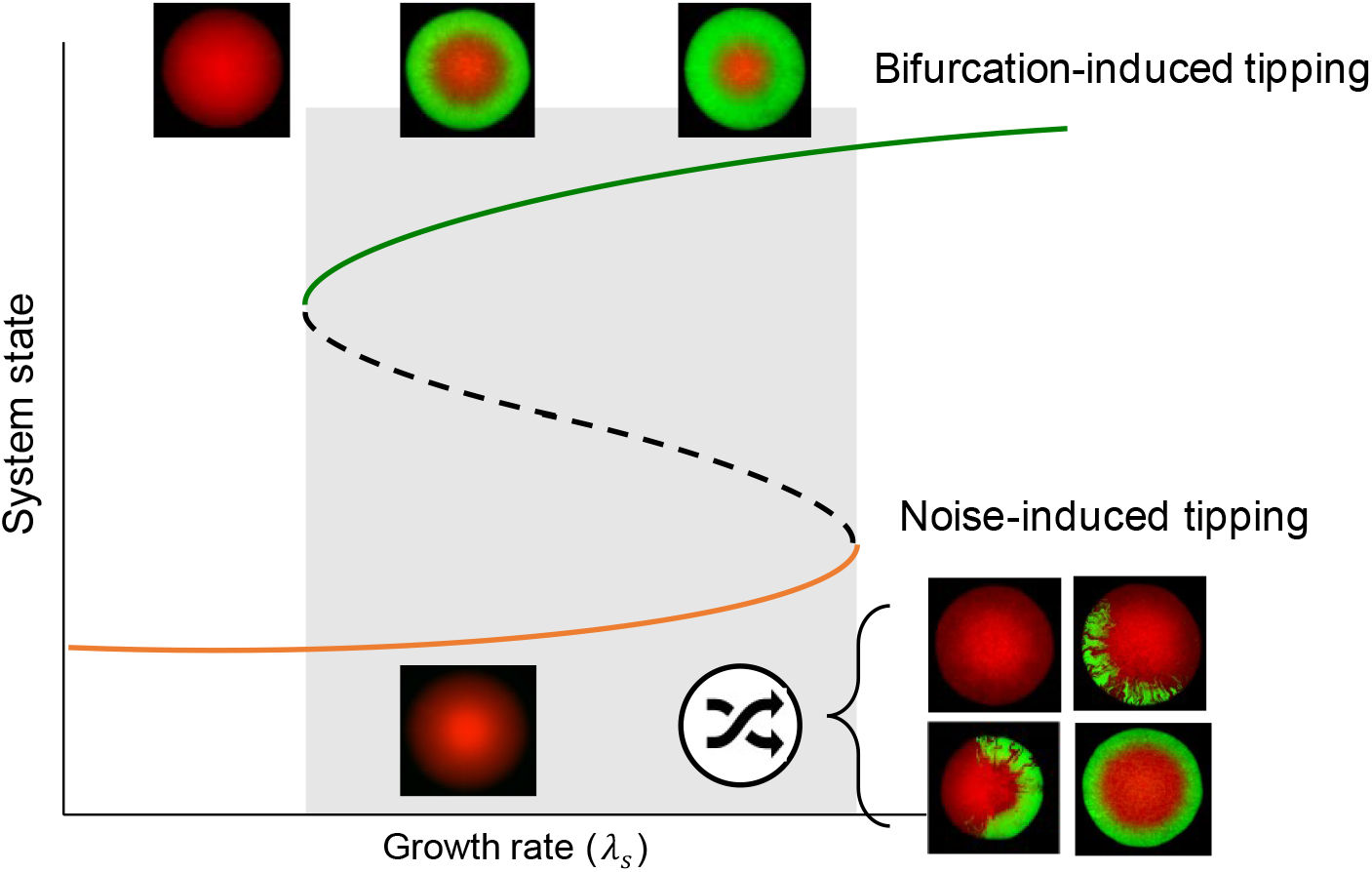
Pattern multistability emerges from a bistable system. This figure illustrates how diverse colony patterns emerge from a bistable system, starting from a single cell under varying external conditions, specifically the maximal growth rate provided by different substrates. When a single bacterial cell carrying a bistable switch begins to grow from a green state within the bistable range, a ring-like pattern consistently forms. Decreasing the substrate nutrient quality (modifying the growth rate *λ_s_*) results in a smaller ring width until the ring eventually disappears (upper row). This ring-like pattern formation can be explained by a deterministic bifurcation process, triggered by decreased cell growth rates in the inner and top regions of the colony during establishment (Fig. 2a). When a single cell begins to grow from an initial red state, where a spontaneous state shift to green may occur due to noise-induced tipping, the spatial system allows for alternative patterns to emerge (bottom row). Range expansion amplifies cellular fluctuations, leading to pattern diversity through the founder effect at the expanding front (Fig. 4a). By controlling the initial state of the single cell, colony patterns with varying degrees of asymmetry can be generated.

The spatial expansion of the bacterial colony spontaneously generated heterogenous structures within the microenvironment (e.g., nutrient access), which enabled position-dependent bifurcation (critical transition) of cell fate and the formation of ring-like patterns (the top row in Fig. 5). The bifurcation tipping point, which defines the boundary of green and red cells, depended on both the intrinsic regulatory properties (e.g. the critical transition growth rate *λ_c_*) and the external growth conditions (*λ_s_*). This deformation of fate landscape is a consequence of global gene regulation in response to growth variations, differing from classic morphogenesis that typically requires a specific gradient of signalling molecules.^44–46^

We also elaborated that range expansion amplifies the cellular fluctuations to pattern diversity through the founder effect at the expanding front (the bottom row in Fig. 5). By controlling the initial state of the single cell, we were able to generate colony patterns with different degrees of asymmetry. Without complicated regulation and prepattern information, such a noise-induced symmetry-breaking mechanism may be relevant in other prokaryotic and eukaryotic systems and could underpin multicellular development and organization.

## Acknowledgements

This work was funded by the National Key Research and Development Program of China (Grant No. 2018YFA0903400), the National Natural Science Foundation of China (Grant No. 32071417 and Grant No. 32301225), the Strategic Priority Research Program of the Chinese Academy of Sciences (Grant No. XDB0480000), the Guangdong Basic and Applied Basic Research Foundation (Grant No. 2021A1515110863), and the Shenzhen Science and Technology Program (Grant No. ZDSYS20220606100606013). We extend our gratitude to Prof. Wei Huang, Prof. Shuqiang Huang, Prof. Hui Sun, and Mr. Yuejian Mo for their technical support. We extend our gratitude to Prof. Xuefei Li, Dr. Nan Luo, and the members of the Fu lab for their enriching discussions. Additionally, we acknowledge the Shenzhen Infrastructure for Synthetic Biology for their invaluable instrument support and technical guidance.

## Author Contributions

X.F. was responsible for conceptualization, supervision, funding acquisition, writing, editing, and project administration. The experimental study was conceived by P.C., J.Z., and X.F.. P.C, J.Z., X.F. designed experiments. P.C., J.Z., and Z.M. performed experiments. P.C designed the numerical study and executed simulations under the supervision of X.F.. P.C. and J.Z., under the supervision of X.F., designed and performed the data analysis pipeline. The manuscript was jointly authored by P.C., J.Z., and X.F..

## Methods and Materials

### Bacterial strains and plasmids

All experiments utilized strains derived from the *E. coli* K-12 NCM3722 background, with strain NH3 specifically generated by the deletion of the *fliC* gene and *lac* operator. The wild-type strain NCM3722 was generously provided by Dr. Chenli Liu. All mutual repression circuits were obtained from the pECJ3 plasmid (gift from Dr. James Collins, Addgene plasmid # 75465)^21^ and harbored in a plasmid with a ColE1 origin. The sequence contents of genetic parts are provided in Table S8.

### Growth medium

All batch culture and colony growth experiments were performed using MOPS-buffered defined medium^47^ supplemented with appropriate antibiotics. The MOPS buffer contains 40 mM MOPS (Sigma-Aldrich V900306), 4 mM Tricine (Sigma-Aldrich V900412, adjusted to pH 7.4 with KOH), 0.01 mM FeSO_4_, 0.276 mM K_2_SO_4_, 0.5 μM CaCl_2_, 0.525 mM MgCl_2_, 50 mM NaCl, 1.32 mM K_2_HPO_4_ and micronutrient mixtures (3 nM (NH_4_)_6_Mo_7_O_24_, 0.4 μM H_3_BO_3_, 30 nM CoCl_2_, 10 nM CuSO_4_, 80 nM MnCl_2_, 10 nM ZnCl_2_). Unless otherwise specified, our MOPS buffer contains 9.5 mM NH_4_Cl as nitrogen source. Different growth rates were achieved by providing different compositions of nutrients including carbon and nitrogen sources. Detailed information on these media is provided in Table S4. The chemical inducers, IPTG (Sigma-Aldrich I6758, isopropyl β-D-1-thiogalactopyranoside) and cTc (Aladdin C103023, Chlorotetracycline hydrochloride), were diluted to concentrations of 0.2 mM and 10 ng/mL, respectively, when required to induce the transition of cells to the red and green states.

The bacto-agar (BD, 214010) was added to the growth medium, and the agar concentration was 1.5% (w/v) unless otherwise specified. A total of 60 mL molten agar gel was poured into a tissue culture plate (JET BIOFIL, TCP011001, 118.4mm×82.48mm×11.77mm) or 35 mL of the agar gel was poured into a petri dish (9 cm diameter, JET BIOFIL, TCD000090) to a final thickness of approximately 6 mm. The agar was allowed to harden at room temperature for 90 minutes before using.

### Batch culture growth and growth rate measurement

Unless otherwise specified, cell culture and growth rate measurements were conducted in three steps: seed culture, pre-culture, and experimental culture. Initially, strains were streaked on LB agar plates from glycerol stocks stored at -80°C. The plates were then incubated at 37°C for 10-12 hours. Subsequently, a single colony was selected from the agar plate and inoculated into a 14 mL tube containing 3 mL LB medium supplemented with an appropriate antibiotic. The culture was agitated in a shaker (220 rpm, 37°C, Shanghai Zhichu Instrument) overnight to serve as the seed culture. During the pre-culture step, cells were transferred to a 0.22 µm filter and washed with pre-warmed experimental medium, ensuring that remnants of the LB medium were removed. Washed cells were then diluted into the experimental medium, with an initial OD_600_ (optical density at 600 nm) of approximately 0.01. Successive dilutions were performed once the OD_600_ reached 0.2, and this process was repeated for several rounds to maintain a balanced growth condition. Samples for further experiments or quantitative measurements were collected when the OD_600_ was less than 0.2.

For the growth rate measurement, cells were maintained in the pre-culture steps for at least 10 generations to establish steady-state growth. Experimental cultures were initiated by diluting the pre-culture to an OD_600_ of approximately 0.02. Measurements were taken at five to eight points within a range of OD_600_ from 0.03 to 0.2. The growth rate (λ) was calculated by fitting the data to an exponential growth curve.

Unless otherwise stated, the pre-culture and experimental culture steps were carried out in a water-bath shaker (150 rpm, 37°C, Shanghai Zhichu Instrument) using a 29 mm × 115 mm test tube with no more than 10 mL of medium.

### Single colony growth in agar plates

Seed cultures and pre-cultures were prepared following the previously described protocol. Once the experimental culture reached approximately 0.1 OD_600_, an aliquot of the culture was taken and diluted into MOPS buffer. For cultures grown in rich medium, the dilution was adjusted to an OD_600_ of 1E-6, while cells growing in minimal media were diluted to an OD_600_ of 5E-7. Before spreading the cells, agar plates were allowed to dry in a hood for 20 minutes. Subsequently, 100 μL of the diluted cell culture was plated onto the agar plates and spread using a glass spreader to achieve approximately 30 colonies on each plate (see Extended Data Fig. 1b).

For the FACS protocol (see Extended Data Fig. 1d), the cell culture was diluted with MOPS buffer to an OD_600_ of approximately 0.001 to maintain the event number under 100 events/s at a flow speed of 10 μL/min. Subsequently, a single cell was inoculated onto the agar plates, ensuring 9 mm between two adjacent cells, resembling the well pattern of a 96-well plate.

The plates were placed in an upright position in a water-bathed incubator (Shanghai Yiheng, GHP-9160) set at 37 ℃, with a humidity of 60% to prevent water loss from the agar. For time-lapse imaging of colony growth dynamics, similar growth conditions were maintained using a custom-made hood on the microscope (Nikon, Ti-E).

### Flow cytometry analysis

The fluorescence intensity distribution of each sample was measured using a flow cytometer (CytoFLEX S, Beckman Coulter Life Sciences) equipped with excitation/emission filters of 488 nm/525 nm (FITC) for GFP and 561 nm/610 nm (ECD) for mCherry. The instrument was operated using CytExpert 2.5 software. Gain settings for FSC, SSC, FITC, and ECD channels were adjusted to 250, 250, 250, and 1500, respectively. The trigger level was manually set on SSC-H based on different conditions. The flow rate was maintained at 60 µL/min, and at least 50,000 events were recorded per sample. To ensure accurate measurements, the detected event number per second was kept below 3000 events/s.

### Fluorescence-activated cell sorting (FACS)

The cell sorter (CytoFLEX SRT, Beckman Coulter Life Sciences, Cyt Expert SRT 1.1 software) was used. The gain for FSC, SSC, FITC and ECD channels were set to 250, 250, 250 and 3000, respectively. The trigger level was manually set on SSC-H to 5000.

For the single cell sorting experiment, we extracted an aliquot of the experimental culture and diluted it with MOPS buffer to achieve an OD_600_ of approximately 0.001. This was done to maintain the event number below 100 events/s at a flow speed of 10 μL/min. Subsequently, a single cell was sorted onto agar plates or well plates using the standard 96-well plate mode, ensuring 9 mm between two adjacent cells. The straight down stream mode was selected for the single cell sorting process.

### Sorting of cells from colony

The entire colony was eluted into 2 mL of MOPS buffer and washed multiple times to ensure retrieval of all cells within the colony. Before and during the sorting experiment, the samples were maintained at 4 ℃. Different subpopulations of cells were sorted according to their fluorescence intensities (Extended Data Fig. 5c and Supplementary Fig. 2) into 5 mL round-bottom test tubes (Falcon, 352052) and 15 mL centrifuge tubes (Corning, 430791) containing 1 mL of RNAProtect (Qiagen, 76506). The sorting rate was kept below 9000 events/s to minimize the false positive ratio and the flow rate was maintained at approximately 25-35 µL/min. Approximately 1 million cells were sorted into tubes for each subpopulation. The sorted cells were then mixed by vortexing for 5 seconds and incubated for 15 minutes at room temperature.

### Bacterial RNA extraction and library construction

We adopted the previously reported mothed MiniBac-seq^48^ to prepare the RNA-seq library with ultra-low amount of RNA. Following cell sorting, the samples underwent centrifugation at 5000g for 15 minutes at 4 ℃ to remove the supernatant, leaving approximately 1 mL of residual volume for each sample. Subsequently, the cells were resuspended and transferred to 1.5 mL microcentrifuge tubes. Another round of centrifugation at 5000g for 15 minutes at 4 ℃ was performed to discard the remaining supernatant. The cell pellet was resuspended in 50 µL TE buffer (10 mM Tris-Cl, 1mM EDTA, pH 8.0, RNase-free, Invitrogen, AM9849-500mL), supplemented with 12.5 mg/mL lysozyme (Sigma-Aldrich, 62970-5G-F) and 16.7% (v:v) Proteinase K (Qiagen, 19131, 2 mL). The suspension was then incubated at 37 ℃ for 15 minutes at 500 r.p.m.

The bacterial RNA was extracted using 100 µL RNAClean XP beads (Beckman, A63987, 40mL). Following purification, the rRNA and non-coding ssrA were eliminated using a customized ssDNA library (refer to [33]). Subsequently, rRNA-free total RNAs underwent an additional purification using RNA cleanup XP beads. The reverse transcription for first-strand synthesis was then conducted using Maxima H Minus Reverse Transcriptase (Thermo Fisher, EP0753, 40000 units, 200 U/µL). The resulting cDNA was purified once more, this time using AMPure XP beads (Beckman, A63881, 60mL). Finally, the second strand synthesis was carried out utilizing the NEBNext Second Strand Synthesis Kit (New England Biolabs, E6111S).

The concentration of the purified double-strand cDNA was quantified using Qubit dsDNA HS Assay Kits (Thermo Fisher Scientific, Q32854) on a Qubit 4 Fluorometer (Invitrogen). Subsequently, the cDNA underwent tagmentation with N7 adaptor-incubating transposase in TruePrep Homo-N7 DNA Library Prep Kit for Illumina (Vazyme, TD512 or TD513 depending on the total cDNA abundance of different samples). This was followed by PCR amplification for approximately 9-15 cycles according to the initial concentrations of DNA. PCR also introduced specific i5 (NEB, 7780) and N8 (Vazyme, TD203) index to the two ends of the product. The oligonucleotides used in the library preparation were listed in Table S10. The resulting products were then quantified using the Qubit 4 Fluorometer, and their size distribution was analyzed using High Sensitivity D1000 Reagents (Agilent, 5067-5585) on an Agilent TapeStation 4150. The libraries that passed the quality assessment were subjected to sequencing by Sangon Biotech using the Illumina NovaSeq 6000 system, employing a 2*150 pair-end configurations. Each sample yielded approximately 25 million paired end reads.

### mRNA sequencing and data processing

Sequencing raw data was processed using customed python (3.7.1) scripts (https://github.com/MinTTT/RNA_seq_pip). Briefly, raw sequencing files were trimmed adapters using cutadapt (v4.2)^49^. Next, the sequencing data were aligned to reference sequences using Bowtie2 (v2.4.1)^50^. The referencing sequences were generated according to the sample type, where the genome is NCM3722 (GenBank: CP011495.1). After alignment, reads coverage over the whole genome was calculated. Specifically, all properly mapped reads were separated according to their mapped strands using samtools (v1.9)^51^. mRNA abundance 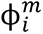 was calculated by followed equation:

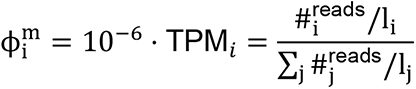

where 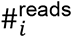 represents the number of mapped reads of gene *i*, *l_i_* denotes the length of gene *i*.

### Mother machine experiment

To prepare the PDMS for the mother machine chips, silicone elastomer (Sylgard 184) was thoroughly mixed with the curing agent in a 10:1 ratio, defoamed, and then poured onto the silicon wafer. The mixture was placed under vacuum (-0.8 kg/cm^2^) for degassing for 10 minutes, followed by removing air bubbles on the surface of the PDMS, and then cured at 80 ℃ for at least 30 minutes. The PDMS was then removed from the silicon wafer. Individual mother machine chips were cut out of the PDMS, and holes were created for the inlets and outlets using a puncher (0.7 mm). Cleaned coverslips (0.13-0.16 mm) were bonded to the feature side of the mother-machine chips using oxygen plasma for 3-5 minutes followed by incubation at 80 ℃ for at least 10 minutes to reinforce the bonding.

Prior to loading the cells onto a machine chip, cells were induced to the red state using 0.2 mM IPTG and cultured to an OD_600_ of approximately 0.6. Cells were then centrifuged and concentrated 100 to 400-fold before being loaded into a mother machine chip, followed by centrifugation at 2500 g for 10 minutes to load individual cells into the side channels. The culture media was initially pumped at a high rate for one hour to clear the inlets and outlets. Subsequently, the media was pumped at 10 μL/min for the duration of the experiment. Cells were allowed to adapt to the environment for several hours before imaging commenced. Pluronic F-108 (Sigma Aldrich, 542342-250G) was added to all media used in the mother machine experiment to achieve a final concentration of 0.85 g/L.

### Mother machine data processing: segmentation and single cell tracking

Customized image processing scripts based on deep learning algorithms^52^ have been developed to analyze mother machine data. This process can be summarized in four main steps: 1. Time-lapse images from each field-of-view (FOV) are aligned to eliminate XY errors during stage movement. A pre-trained model for side-channel detection was then applied to identify channels within the FOVs; 2. Individual cells are segmented using the model for cell detection. The edge details of cells are determined using Otsu’s thresholding algorithm; 3. The midlines of cells are calculated through interpolation to provide an initial guess for cell size determination. Then, the cell size information, including cell mask, length, width, and area, is calculated based on the cell coordinate system^53^; 4. The fluorescent protein expression levels are extracted from images of fluorescence channels. The mask of cells is applied to images, and the median pixel value within channels is used as the fluorescence background. The fluorescence of a cell is calculated by subtracting the fluorescence background from the median value of the pixels belonging to the cell.

### Two-photon microscopy

The Nikon AX-FN microscope, equipped with a 10x air objective (Plan Apo λD NA 0.45), was used to capture images. The resonant mode was used to ensure fast acquisition, with line averaging set to 16x. The scanning resolution was configured at 1024x1024. Excitation wavelengths of 960 nm for the GFP channel and 1040 nm for the RFP channel were utilized. Both fluorescence channels were simultaneously acquired. A Z-step of 5 μm was employed for 3D sectioning. In terms of sample preparation, samples were stored at 4 ℃ for 18-24 hours to allow for fluorescence protein maturation. Throughout the experiment the plates were kept upright.

### Wide-field microscopy

Microscopic images of the bottom view of the colonies were captured using Nikon TiE microscope. Our microscope was equipped with a SpectraX (Lumencor) as epi-illuminator, a 4x objective (Nikon, Flu Plan NA 0.13, for colony acquisition), a 60x objective (Nikon, Plan Apo λ NA 1.4, for single cell acquisition in the agar pad), a 100x objective (Nikon, Plan Apo λ NA 1.5 for single cell tracking in the mother machine), and an ORCA flash4.0 sCOMS camera (Hamamatsu). For fluorescence channels, the dual-channels filter set (Chroma, 59022) was used to capture the GFP and RFP channels.

To characterize the dynamics of the colony expansion or single cells, environmental temperature was controlled by a tailor-made hood to maintain a stable 37 ℃ growth temperature, with a humidity of 60% to prevent agar evaporation.

In the experiments of colony expansion, after spreading the cells, the plate was placed on the microscope stage, and image acquisition began when the colony radius reached approximately 200 µm. The RFP, GFP, and phase channels were captured sequentially, with a frequency of every 15 minutes.

In single-cell tracing experiments, which include both the mother machine and the agar pad setups, the phase channel was captured every 3 minutes, while the RFP and GFP channels were acquired every 9 minutes.

For characterizing colony bottom patterns, the plates were stored at 4 ℃ for 18-24 hours before image acquisition.

### Computational model

The chemo-mechanical model from ref^32,33^, incorporating the mutual repression network, was employed to describe and analyze the spatiotemporal establishment of the ring-like pattern in bacterial colonies. (Supplementary Note 1) The agent-based model adapted from ref^23^, which incorporates the biochemical network with stochasticity (Supplementary Note 2), was utilized to describe the pattern establishment from a red state single cell during the early stage (< 6 hours) of colony formation.

### Colony statistics

We apply the spatial entropy *H*_s_introduced in previous work^54^ to quantify the spatial heterogeneity of the bacterial colonies. Bands with a width of 60 pixels, equivalent to 97.5 µm, positioned 300 to 360 pixels away from the colony centre, were utilized for this analysis (Fig. S16). The spatial entropy can be defined as follows:

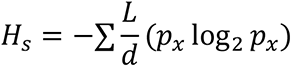

where *p_x_* corresponds to the proportion of all red state pixels to the total pixels and the proportion of green state pixels, L represents the sum of the edge lengths between the red state sectors and the green state sectors, and d denotes the Euclidean distance between the centroids of the red state sectors and those of the green state sectors. To prevent null values of d in spatial entropy calculation (i.e., when the centroids of the red and green states coincide), a relatively small constant (1 pixels) was introduced.

## Data availability

Major experimental data supporting the findings of this study are available within the main text and Supplementary Information.

Sequencing data have been deposited to the National Microbiology Data Center (NMDC10018961).

Flow cytometry data is available at Zenodo (10.5281/zenodo.11817798).

The scripts for mother machine data analysis are deposited to the GitHub (MinTTT/MoMa_process).

Numerical simulation scripts are deposited to the GitHub (MinTTT/Scripts_for_colony_pattern_development).

---

**Extended Data Fig. 1.**
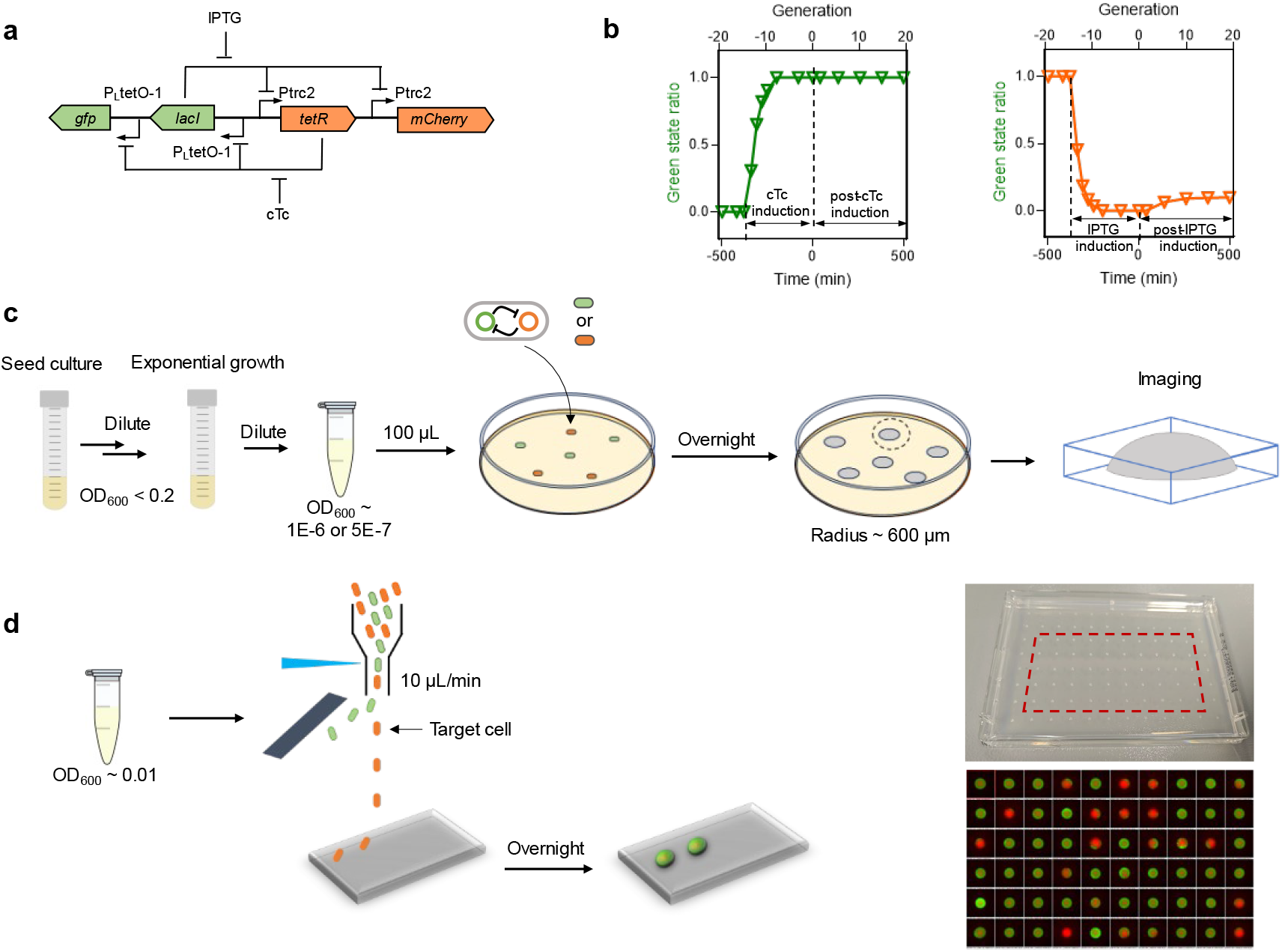
Bistable switch characterization and experimental workflow for single-cell colony culture. (a-b) Schematic illustration of the bistable switch design and demonstration of bistablility. (a) The circuit consists of reciprocal transcriptional repression by TetR and LacI. mCherry and GFP serve as reporters for the TetR high state and LacI high state, respectively. The addition of cTc relieves TetR repression, allowing high expression of LacI and GFP (green state), while induction with IPTG relieves LacI repression, allowing high expression of TetR and mCherry (red state). (b) During exponential growth in RDM glucose medium, following prolonged culture via successive dilution, hysteresis of cellular states is demonstrated. A small population (∼10%) of post-IPTG induction red state cells spontaneously switched to the green state (high GFP state) within 20 generations. (c-d) The experimental workflow for single-cell colony culture: (c) Colonies were spread using a glass spreader to achieve approximately 30 colonies per plate; (d) Single-cell sorting was performed using FACS in standard 96-well plate mode (see Methods). Sixty colonies were selected for further analysis on each plate (right panel).

**Extended Data Fig. 2.**
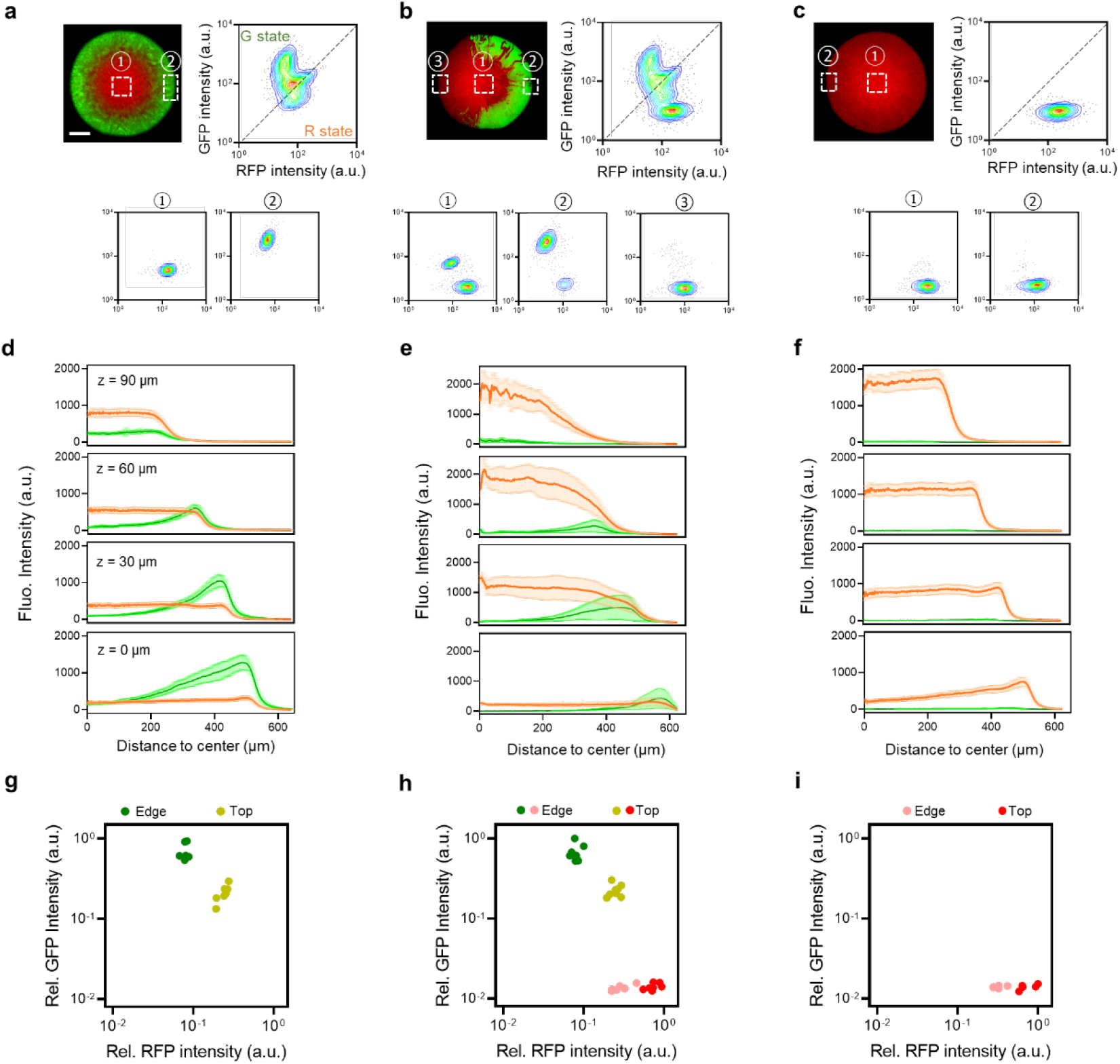
Characteristics of three typic types of patterns. (a-c) Flow cytometric analysis of whole individual colonies (upper panel) representing each pattern type, with different regions of each colony analyzed (bottom panel). The regions analyzed include the colony top (①) and the colony periphery (② and ③). The top regions consistently maintain the red state, regardless of the peripheral regions’ state. Scale bars indicate 200 µm. (d-f) Fluorescence intensity profiles obtained by two-photon microscopy for the colonies shown in panels (a-c) and Fig. 1b at four selected heights from 0 to 90 µm. RFP intensity increases with height in all three pattern types, while GFP is predominantly distributed at the bottom part of the peripheral regions in the ring-like (a) and hybrid patterns (b). Despite the impact of cell density at different radial and vertical positions on fluorescence intensity, flow cytometric analysis in (c) reveals significant variations in RFP fluorescence intensity at the top or periphery, even though red state cells were present throughout the colony. (g-i) Comparison of phenotypic characterizations of cells picked from the edge and top of the colony for each pattern type. Five individual colonies were selected for each pattern type.

**Extended Data Fig. 3.**
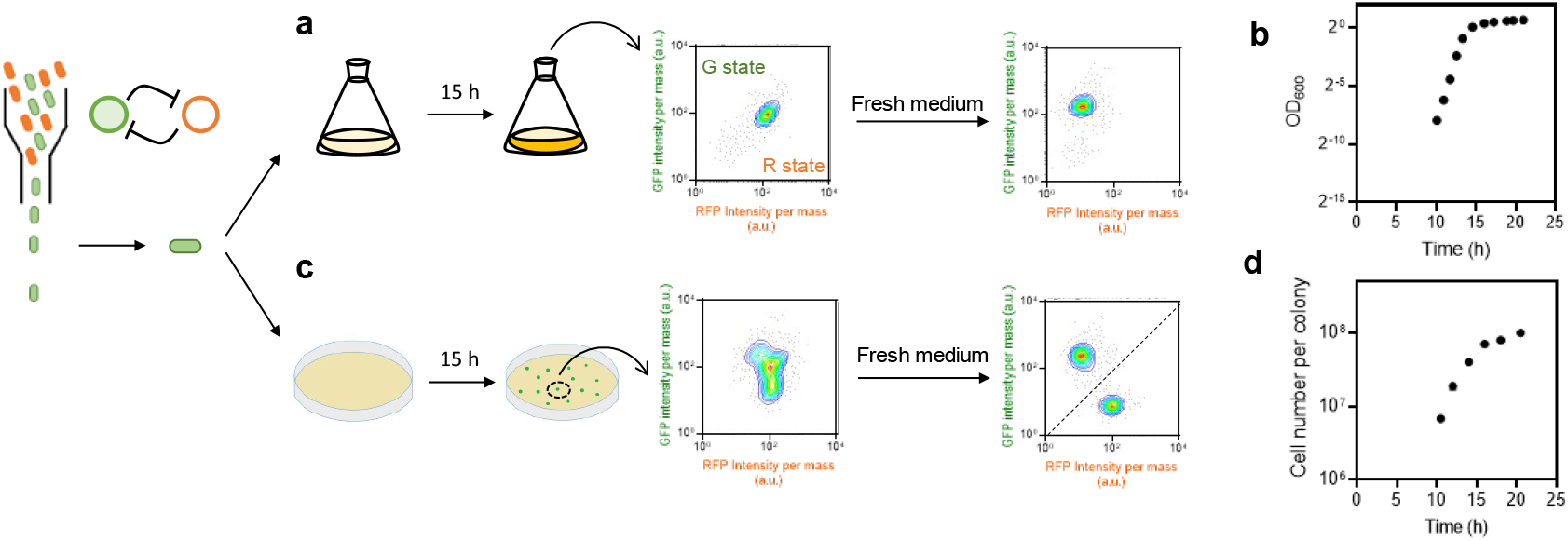
Cell state evolution from a green state single cell in liquid culture and on solid agar. (a) Single green state bacterial cells were introduced into RDM glucose medium by FACS. The culture was maintained under constant shaking to ensure a well-mixed environment. Cell density (OD_600_, panel b) and phenotypic changes were monitored over a set period. Co-activation of GFP and RFP was identified after 15 hours of culture. Subsequent re-culturing of these cells into fresh medium resulted in a homogeneous green-state population. (c) Single green state bacterial cells were inoculated on a solid agar (1.5% w/v) plate supplemented with RDM medium. Colonies from different incubation times were observed for cell density (panel d) and phenotypic changes (Extended Data Fig. 4). The solid medium created a heterogeneous microenvironment, resulting in varied nutrient access and local cell density, significantly influencing the distribution of cell states. Re-culturing cells from a colony grown for 15 hours into fresh RDM medium revealed two distinct phenotypes.

**Extended Data Fig. 4:**
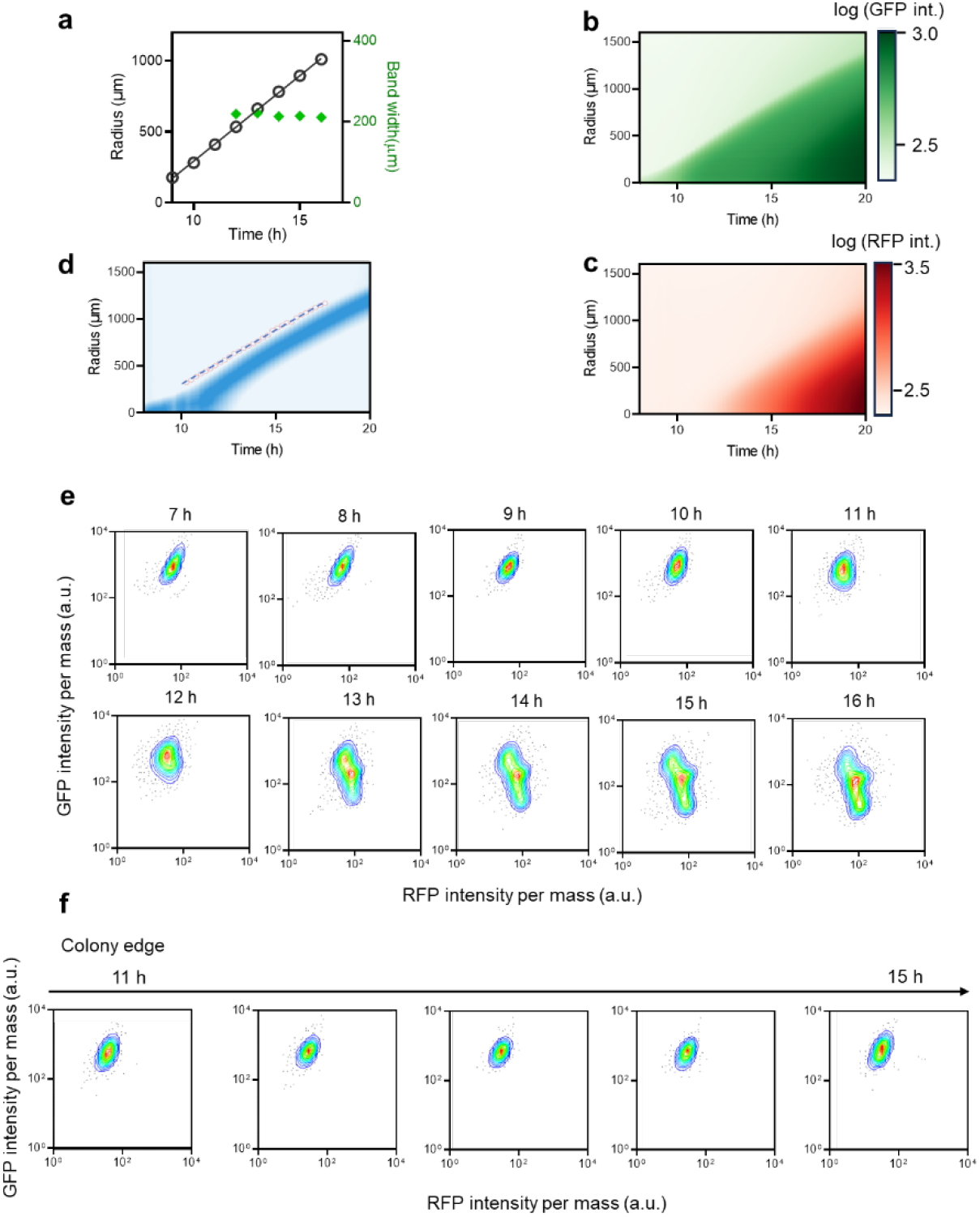
Experimental observations regarding the spatiotemporal establishment of the colony and the evolution of phenotypes. (a-d) Time-lapse expansion kinetics of the bacterial colony were monitored using a wide-field microscope as it grew on a 1.5% agar plate supplemented with RDM glucose medium. (a) The growing colony exhibited linear expansion, with a radial expansion rate of 120.1 µm/h observed from 9 to 16 hours after initial seeding. A constant band width of approximately 216 µm was characterized (green diamonds and panel (d)). (b-c) The radial profiles of GFP intensity (b) and RFP intensity (c) at different times were recorded. Panel (d) describes the changes in the ratio of GFP intensity to RFP intensity over time, characterizing a constant band where GFP expression predominates. (e-f) The time evolution of cell phenotype distribution in entire bacterial colonies (e) and in cells picked from the colony periphery (f) at different times was analyzed. A uniform green state population was present within the initial 10 hours following cell seeding. As the culture time progressed, a transient transition towards a heterogeneous population was observed. A portion of the cells gradually transitioned to the red state as the colony continued to expand both radially and vertically.

**Extended Data Fig. 5:**
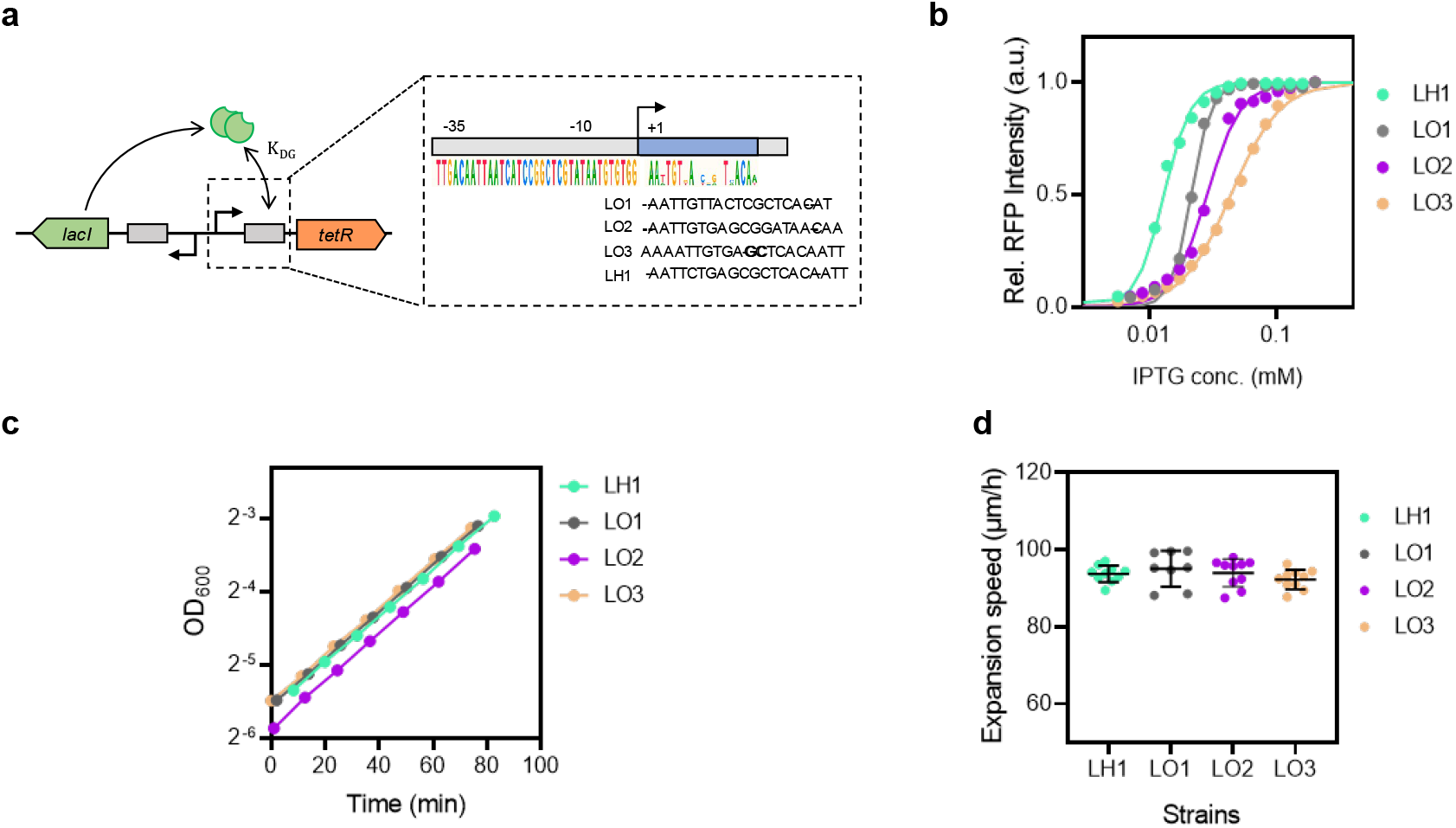
Modification of the critical growth rate *λ_c_* of toggle switch strains. (a) Four representatives of synthetic binding sites for the transcription factor LacI were studied to vary the repression threshold *K_DG_*. (b) IPTG induction response curves of the four studied mutants of the toggle switch indicate the relative values of the repression threshold *K_DG_* for each circuit. Symbols represent experimental data, while solid lines depict parameter fitting lines using the Hill function. A larger effective threshold of the fitted curve indicates a higher repression threshold, following the trend LH1 > LO1 > LO2 > LO3. (c) Exponential growth of different strains measured in MOPS CAA glucose medium. (d) Radial range expansion rates of different strains on a 1.5% agar plate supplemented with MOPS CAA glucose medium. No significant differences were observed.

**Extended Data Fig. 6:**
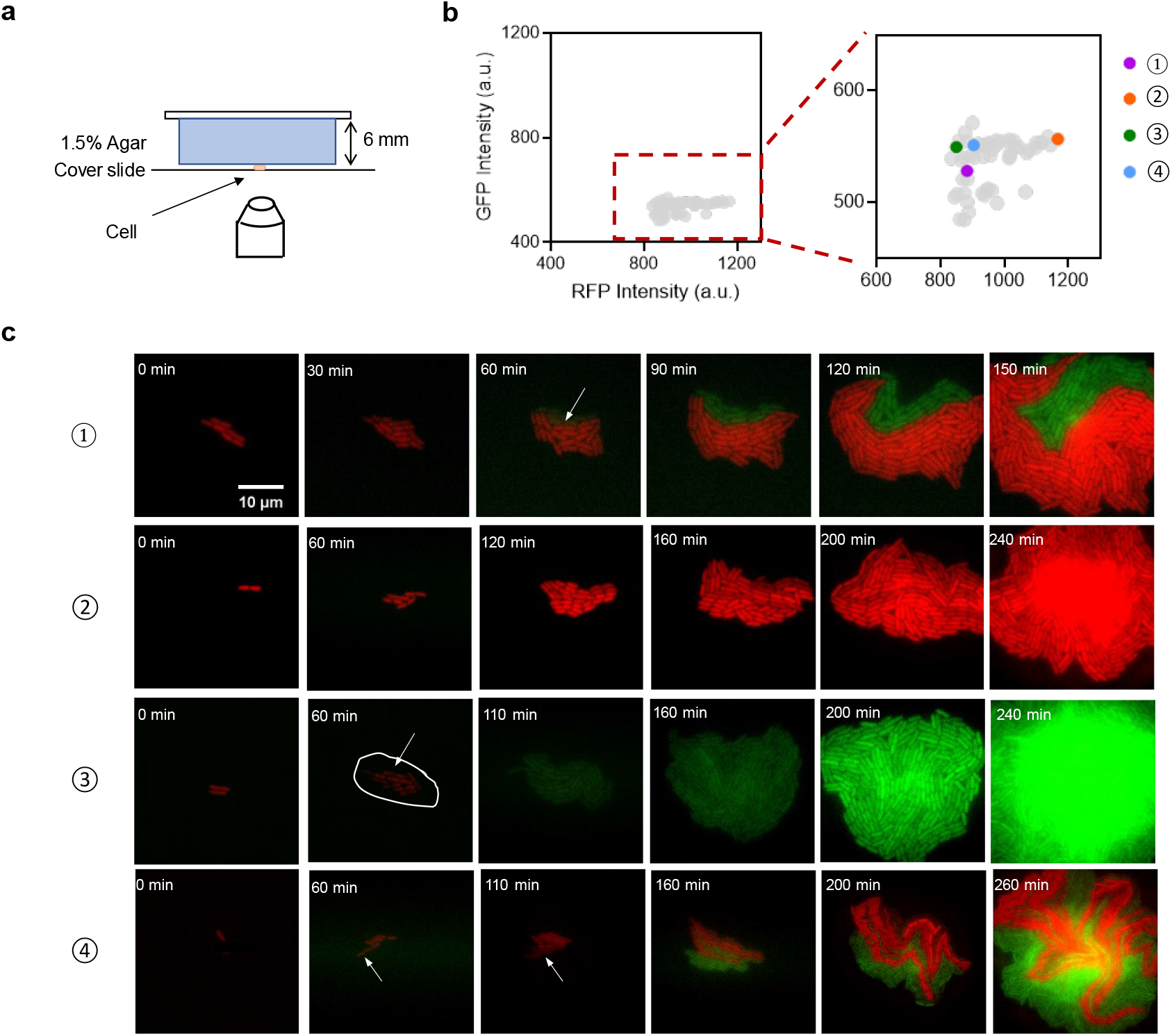
Snapshots of initial red state cells growing on agar pad. (a) Illustration of the agar pad used to monitor the early stages of bacterial colony formation from a single red state cell (see Methods). (b) Fluorescence intensity distribution of cells’ initial state. A total of 55 red state single cells were plotted. The initial conditions of four representative progenitor cells presented in panel (c) are provided. (c) Four representative snapshots of pattern formation from a single cell to a cluster by time-lapse microscopy. The scale bar indicates 10 µm. The initial condition with relatively high RFP intensity (②) results in a homogeneous red state pattern after four hours, while the cell with relatively low RFP intensity (③) undergoes a spontaneous state transition to the green state within 110 minutes, resulting in a homogeneous green cluster. Cells originating from ① and ④ produce mixed clusters containing both red state and green state cells.

**Extended Data Fig. 7:**
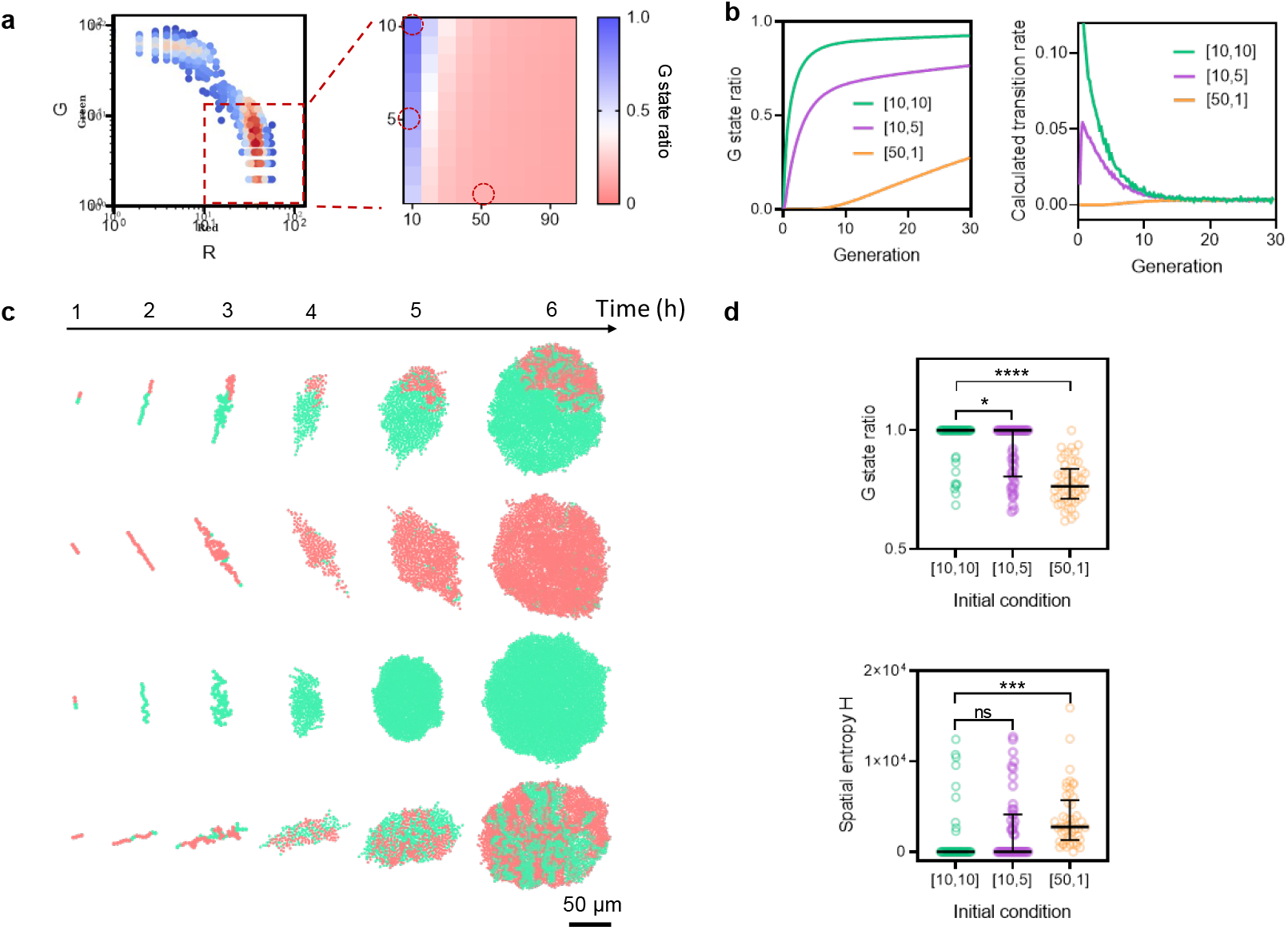
Simulations reveal that the state transition rate during the early stage of pattern formation depends on the initial condition of gene expression. (a) The G state ratio after 30 generations of growth, dependent on the initial condition. ’R’ represents TetR and RFP expression, while ’G’ represents LacI and GFP expression. Simulations were conducted with 200,000 individual cells for each condition. (b) Evolution of the G state ratio and state transition rate over time for three selected initial conditions. The state transition rate is monitored by counting the ratio of cells that switched to the G state within a specified time interval. The transition rate stabilizes at a constant value after approximately 15 generations post-seeding. (c) Four representative snapshots illustrating pattern formation from a single cell to a cluster, observed via a stochastic agent-based model (see Supplementary Note 2). Light red denotes the R state, while spring green represents the G state. (d) The G state ratio and spatial entropy (see Methods) of clusters at 6 hours (as shown in panel c) after the initial seeding of single cells from different initial conditions. Spatial entropy computation treats each individual cell as a mass point, considering its centroid as the position. The plotted lines depict the median and interquartile ranges. To contrast between groups, an unpaired, two-sided t-test with a 95% confidence interval was applied.

**Extended Data Fig. 8:**
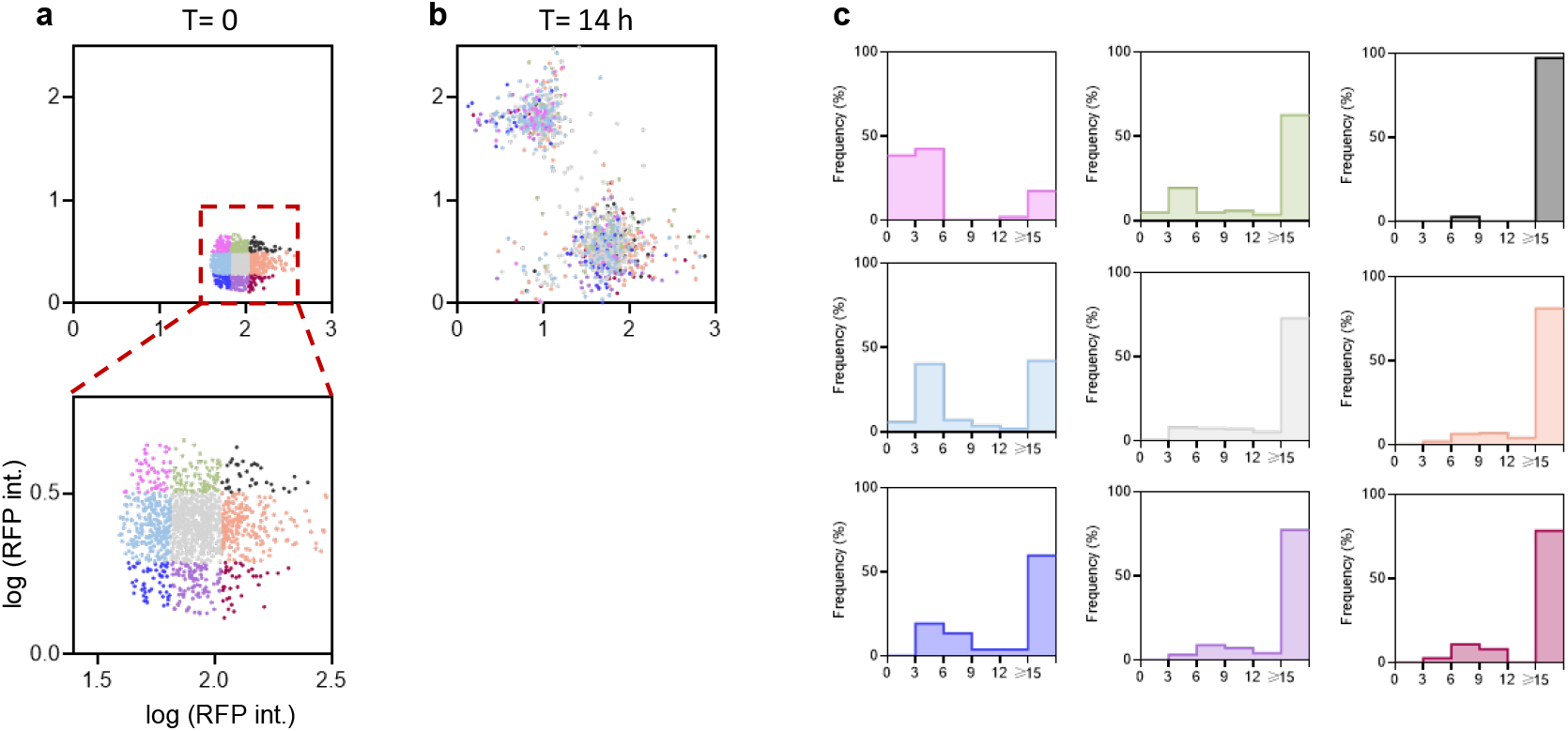
Characterization of spontaneous state transition under a balanced growth condition monitored by the mother machine microfluidic experiment. (a) Scatter distribution of 1383 mother cells, all initially characterized as red state cells. Nine subpopulations are categorized based on their GFP and RFP intensities. (b) Scatter distribution of cells after 14 hours of culture under balanced grow condition in RDM glucose medium. Approximately 36% of cells were observed to switch to the green state. (c) Histograms depicting the state transition time for each subpopulation. Transition times greater than 15 hours indicate that the cells did not change their state throughout the entire experiment. Colors correspond to the initial conditions described in panel (a).

