## Supplementary material for "Colony pattern development of a synthetic bistable switch": Figure S1

### Supplementary Notes

#### Supplementary Note 1: A continuum model for colony expansion.

To understand the interaction between the microenvironment (in this case, nutrients), cell states, and the establishment of a colony. We regard the colony as a growing mass of incompressible fluid. The phase-field method provides a convenient framework for describing the fluid on phenomenology. It introduces an auxiliary phase, $\phi(\mathbf{r}, t)$, that smoothly changes from 0 outside the colony to 1 inside. For simplification of the model, we only consider evolutions in the radius and height of the colony, i.e., the coordinate system is $\mathrm{r}(r, z)$. The schematic of this model is illustrated in **Fig. S1.**

##### 1. Phase field model

As previous works have established1-4, the time evolution of $\phi$ can be given by the equation,

$$\frac{\partial \phi}{\partial t} + \mathbf{u} \cdot \nabla \phi = -\Gamma[G'(\phi)/\epsilon - \epsilon \nabla^2 \phi]. \qquad \text{(S1)}$$

Here, the phase field $\phi$ is advected by the velocity field $\mathbf{u}$, $\Gamma$ is a Lagrange multiplier, $G(\phi)$ is a double-well potential function that forces the field $\phi$ to incline to two fixed points 0 and 1, and the last term $\epsilon \nabla^2 \phi$ represents the surface energy, where $\epsilon$ determines the width of the interface between air and colony.

The growth of the cells can be expressed as:

$$\nabla \cdot \mathbf{u} = k_g(\mathbf{r}, t), \qquad \text{(S2)}$$

where $k_g(\mathbf{r}, t)$ refers to the local growth rate which is a function of the dynamics of metabolism of cells, which will be discussed later.

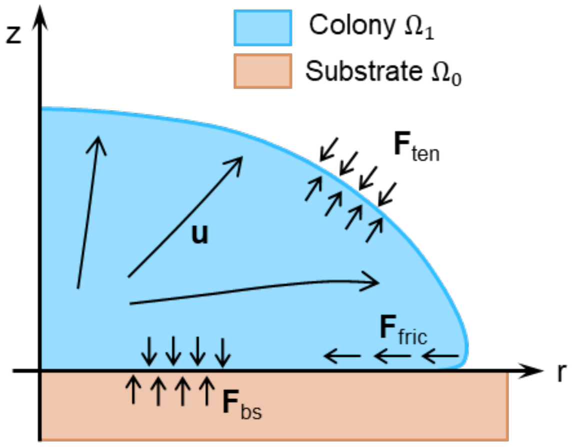

**Fig. S1:** The schematic of the phase field model for bacterial colony development. The blue region indicates the field of a colony which is determined by phase field $\phi$, and the brown portion beneath the colony region is determined by phase filed $\chi$. Cell growth and interfacial forces shape the colony.

The colony is considered as an incompressible fluid flowing through a porous medium (the matrix that consists of *E. coli* cells)5. The velocity field $\mathbf{u}$ can be determined by Darcy's law,

$$-\nu_0\mathbf{u} - \nabla(\phi p) + \mathbf{F}_{ten} + \mathbf{F}_{bs} + \mathbf{F}_{fric} = 0, \quad \text{(S3)}$$

where $\nu_0$ is the viscosity, $p$ is the pressure that originates from the cells' migration and growth. The $\mathbf{F}_{ten}$, $\mathbf{F}_{bs}$, and $\mathbf{F}_{fric}$ are interfacial force terms, which are given as follows:

$\mathbf{F}_{ten}$ denotes the surface tension of the thin liquid layer on the surface of colony. Assuming it is proportional to the colony perimeter 1, we have:

$$\mathcal{H}_{ten} = \gamma L = \gamma/2\int \mathrm{d}\mathbf{r}\{\epsilon(\nabla\phi)^2 + G/\epsilon\}. \quad \text{(S4)}$$

It results in the surface tension $\mathbf{F}_{ten} = -\gamma(\epsilon\Delta\phi - G'(\phi)/\epsilon)\nabla\phi$.

$\mathbf{F}_{bs}$ refers to the dhesion and frication between the colony and the agar substrate. We introduce another phase field2, $\chi(z)$ representing the agar substrate, and it is defined as $\chi(z) = 1/2 - 1/2\tanh(3(z - z_s)/\delta)$, where, $z_s$ is the location of the agar surface, and $\delta$ is the thickness of the interfacial layer between the agar surface and air. The interaction potential between the colony and the agar is then given as,

$$\mathcal{H}_{adh} = \int \mathrm{d}\mathbf{r}\{\phi^2(\phi-2)^2(\underbrace{-2A\delta^{-1}G(\chi)}_{\text{drag}} + \underbrace{g\chi(z+\epsilon)}_{\text{repulsive}})\} \quad \text{(S5)}$$

In this equation, the first two terms $\phi^2(\phi-2)^2$ ensure that the interaction potential only presents in the region of colony, and the last two terms denote the drag and repulsive potential. Here, $A$ is the adhesion energy in the interface of the agar, and $g$ denotes the repulsive potential that prevents the colony from penetrating into the agar. Therefore, the adhesion force between the colony and the agar can be described as, $\mathbf{F}_{adh} = -4\phi(2 - 3\phi + \phi^2)W(\chi)\nabla\phi$. The friction between the colony and the agar originates from the migration of the colony, is defined as $\mathbf{F}_{\mathbf{fric}} = -\eta_s\delta^{-1}\chi\mathbf{u}_{||}$ , where $\eta_s$ denotes the frication coefficient, and $\mathbf{u}_{||}$ is the components of $\mathbf{u}$ in the $r$-direction. As mentioned above, the $\mathbf{F}_{bs}$ is the composition of $\mathbf{F}_{adh}$ and $\mathbf{F}_{fric}$, we have $\mathbf{F}_{\mathbf{bs}} = \mathbf{F}_{\mathbf{bs}} + \mathbf{F}_{\mathbf{fric}}$.

#### 2. Nutrients and cell physiology

The local growth rate $k_g$ is coupled to the nutrients distribution in the colony. We suppose there are two specious of nutrients, $N_1$ and $N_2$, which provide carbon and energy for cell growth. In this study, various media were tested, including minimal media with glycolytic substrates and rich defined media. It was observed that the ring-like band can exhibit regardless of the carbon sources used. The conversion from green to red in cells requires more than 5 generations to transition from one stable state to another. Simulations and experiments demonstrated that a single nutrient can only support cell growth within approximately 50 μm of the nutrient source 6-8. This led to the discovery of a hidden resource pool that supports the growth of cells unable to access fresh nutrients from the colony-agar interface. Given that certain biological processes, such as overflow metabolism9, can provide metabolic byproducts or "waste" from the glycolytic pathway. Our transcriptomic analysis revealed that peripheral cells elevated the glycolytic process and transporter expression(**Fig. S8**), suggesting a nutrient-rich microenvironment and the presence of the overflow metabolism. In contrast, cells at the colony apex show heightened glutamine and lipid metabolism which indicate reallocation of nutrient for balancing building blocks pool. It is hypothesized that cells on the edge of the colony extrude nutrients, thereby bridging gaps in nutrient availability for cells within the colony. we use cross-feeding metabolic processes to describe the dynamics of cell growth, as illustrated in **Fig. S2**.

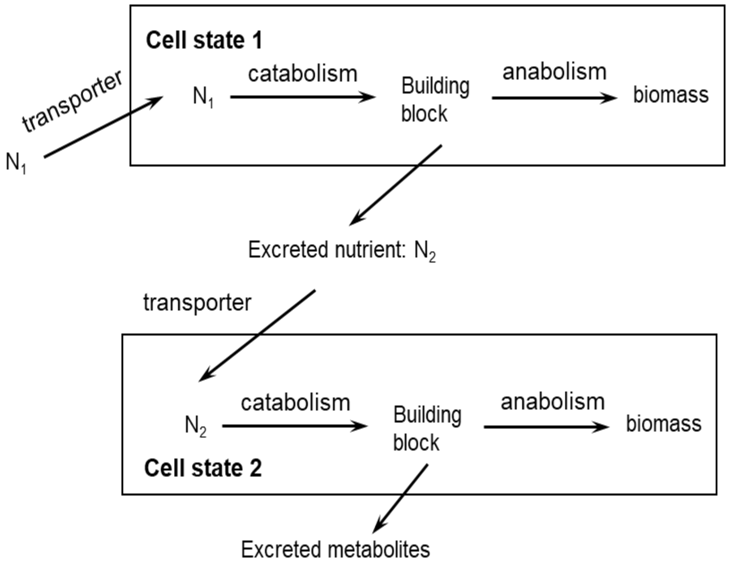

**Fig. S2:** The schematic of the phase field model for cross-feeding metabolic process.

These processes can be summarized in two main steps:

(a) cell growth on $N_1$ and excrete $N_2$.

$$N_1 \rightarrow \text{biomass} + N_2; \tag{S6}$$

(b) cell growth on $N_2$.

$$N_2 \rightarrow \text{biomass} \tag{S7}$$

We use modified Monod equation to approximate the local growth rate. The concentrations of the two nutrients are $C_1$ and $C_2$. We define two Hill equations,

$$\theta_1 = \frac{C_1}{K_1 + C_1}, \quad \theta_2 = \frac{C_2}{K_2 + C_2} \tag{S8}$$

Here, $K_1$, and $K_2$ are the Monod constants corresponding to $N_1$, $N_2$, respectively. We can determine the local growth rate $\lambda(C_1, C_2, \mathbf{r}, t)$ at spatial point $\mathbf{r}$ and time $t$ by,

$$\begin{aligned} \lambda_1 &= \lambda_1^{\max} \cdot \theta_1 \\ \lambda_2 &= \lambda_2^{\max} \cdot \theta_2 \\ k_g &= \lambda_1\theta_1 + \lambda_2(1-\theta_1) \end{aligned} \tag{S9}$$

where, $\lambda_1^{\max}$ and $\lambda_2^{\max}$ are the growth rates of the cells growing in the saturated sole nutrient, respectively. $\lambda_1$ represents the growth rate of cells utilizing the nutrient $N_1$, similarly, $\lambda_2$ corresponds to cells using the nutrient $N_2$. It is noteworthy that $\lambda_1^{\max}$ is identical to $\lambda_s$ ,which represents the global nutrient quality of the substrate media for colony growth in our experimental observations. $k_g$ is the local growth rate that combines contributions from both growth rates, $\lambda_1$ and $\lambda_2$, weighted by the Hill equation $\theta_1$. The dynamics of the concentration of nutrient $N_1$ is given by,

$$\begin{aligned} \partial_t C_1 &= D\nabla^2 C_1 && \text{in } \Omega_0, \\ \partial_t C_1 &= D\nabla^2 C_1 - f_1\lambda_1\theta_1 && \text{in } \Omega_1, \end{aligned} \tag{S10}$$

where, $D$ is the diffusion coefficient, for convenience, we suppose the nutrient diffusion coefficient is a constant in all regions for two nutrients. $f_1$ is the uptake flux of $N_1$.

$$\begin{aligned} \partial_t C_2 &= D\nabla^2 C_2 && \text{in } \Omega_0, \\ \partial_t C_2 &= D\nabla^2 C_2 + p_2\lambda_1\theta_1 - f_2\lambda_2(1-\theta_1) && \text{in } \Omega_1, \end{aligned} \tag{S11}$$

where, $f_2$ is the uptake flux of $N_2$, and $p_2$ is the excretion flux of $N_2$.

The interface conditions on the colony-agar $\Gamma_{01}$ are defined as follows,

$$\frac{\partial C_1^{\Omega_0}}{\partial z} = \frac{\partial C_1^{\Omega_1}}{\partial z}, \quad \frac{\partial C_2^{\Omega_0}}{\partial z} = \frac{\partial C_2^{\Omega_1}}{\partial z} \tag{S12}$$

The boundary conditions for nutrients are:

$$\frac{\partial C_1}{\partial n} = \frac{\partial C_2}{\partial n} = 0 \qquad \text{on } \Gamma_0 \cup \Gamma_1 \tag{S13}$$

The colony and the agar regions, $\Omega_1$ and $\Omega_0$, are defined as $\phi \geq 9e-1$ and $\chi \geq 9e-1$. The interface of the colony and the agar, $\Gamma_{01}$ is defined as the $\max(\phi \cdot \chi)$ along the $z$ direction.

##### 3. The dynamics of the toggle switch

The genetic circuit, comprising two mutually repressed genes, *lacI* and *tetR*, is coupled to the physiological states of the cell. These genes exhibit expression rates that can be approximated as functions of the growth rate10. As previous studies have established, the dynamics of the gene expression rate, in the absence of intrinsic regulatory logic, are described by $\alpha_R(k_g)$ and $\alpha_G(k_g)$. These functions capture how the constitutive expression of TetR and LacI genes responds to changes in the growth rate. The intrinsic repression can be modeled by Hill functions, $H_R([G])$ and $H_G([R])$,

$$\begin{aligned} H_R([G]) &= \tau_R + \frac{1-\tau_R}{1+([G]/K_R)^{n_R}} \\ H_G([R]) &= \tau_G + \frac{1-\tau_G}{1+([R]/K_G)^{n_G}} \end{aligned} \tag{S14}$$

here, $[G]$ is the concentration of LacI, and $[R]$ represents the concentration of TetR.

The overall dynamics of the circuits coupled to phase filed $\phi$ are governed by the following partial differential equations:

$$\begin{aligned} \partial(\phi[G])_t &= -\nabla\cdot(\phi[G]\mathbf{u}) + \alpha_G\phi H_G([R]) \\ \partial(\phi[R])_t &= -\nabla\cdot(\phi[R]\mathbf{u}) + \alpha_R\phi H_R([G]) \end{aligned} \tag{S15}$$

where, the advection terms are coupled with phase filed $\phi$ and the velocity field $\mathbf{u}$ , accounting for the cell movement and protein dilution.

#### 3. Numerical algorithm

To reduce numerical complicity and computational resource usage, the system was simulated within a 2-dimensional field denoted by $\mathbf{r}(r, z)$. Here, $r$ axis denotes the radial expansion space of the colony, and $z$ axis indicates the space that the colony grow vertically. The colony-agar interface is parallel to the $r$ axis and perpendicular to the $z$ axis.

The discrete-continuum model is composed of three main parts: 1. The phase-field part that determines the colony shape and periphery; 2. The reaction-diffusion equations that model nutrients diffusion and consumption, and determine the local growth rate that is coupled to the phase-field part. 3. the dynamics of the toggle switch that is coupled to phase-field part.

The algorithm implemented for the phase-field parts and the dynamics of the toggle is similar to those used in Cao, Y. et al.2, Xiong, L. et al. 3, and Qin, B. et al. 4 As diffusion and consumption are significantly faster than colony expansion rates and the dynamics of the toggle switch, a hybrid scheme was used to simulate the reaction-diffusion equations. Nested grids were created using 3x3 binned grids drawn from the grids of the phase field. In each iteration step, the steady-state solution of the reaction-diffusion equations was calculated using the forward Euler method by time step $\delta_t/N_t$, where $\delta_t$ is the time step used to simulate the dynamics of the phase field and the toggle switch. As the initial values of the nutrient fields $C_1$ and $C_2$ are carried from the previous step, the steady-state solution can be converged rapidly within a few iterations. The parameters used in simulation please refer to Table S9.

#### Supplementary Note 2: An agent-based model for colony establishment.

##### 1. Modeling the toggle switch

Despite previous works indicating that gene expression follows a burst-like mode, we propose that the toggle switch exhibits a simple chemical reaction system where all proteins are produced one by one. We apply the quasi-steady-state assumption to reactions with the Hill function rate, without decomposing them into elementary reactions11. We have four reactions:

$$\begin{aligned} G &\xrightarrow{\alpha_G \cdot H_G(R)} G+1, \quad G \xrightarrow{\lambda \cdot G} G-1 \\ R &\xrightarrow{\alpha_R \cdot H_R(G)} R+1, \quad R \xrightarrow{\lambda \cdot R} R-1 \end{aligned} \tag{S16}$$

where $\alpha_R$, $\alpha_G$ represents gene expression capacity, and $H_G(R)$, $H_R(G)$ are hill functions, as mentioned in Equation (S14), and $\lambda$ is the growth rate determining the dilution rates of proteins. We employ the classical Gillespie algorithm to generate possible trajectories (A typical trajectory is shown in **Fig. S3**). The parameters used in simulation please refer to Table S9.

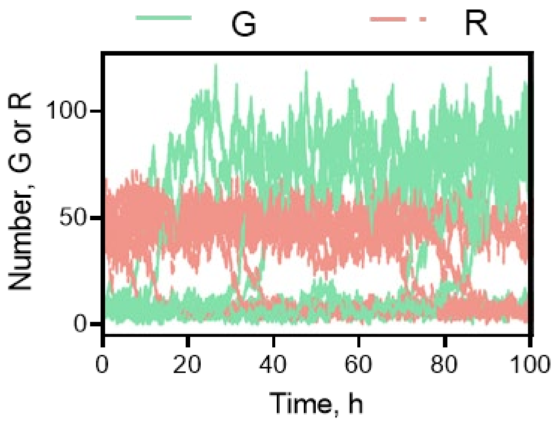

**Fig. S3: A typical trajectory of the toggle switch generated using the Gillespie algorithm.** The green solid line represents the dynamics of the LacI, and the red dashed line indicates the dynamics of the TetR.

##### 2. Modeling colony evolution

This model is based on a 3D colony model previously published in eLife 6. We integrate the modeling scheme into the 3D colony model, the algorithm is shown in **Fig. S4**.

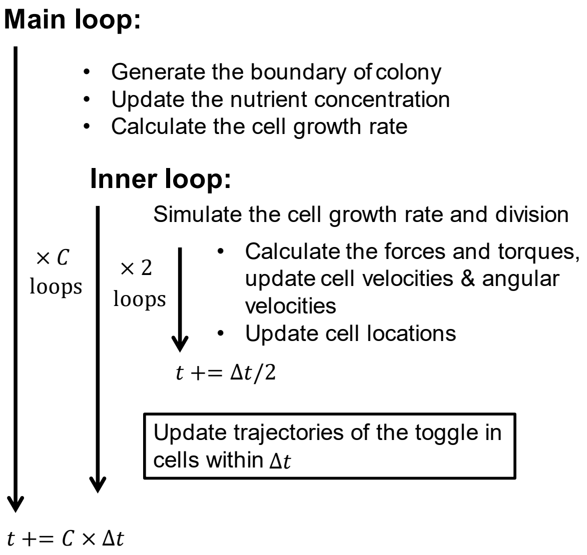

**Fig. S4: the algorithm of the 3D colony incorporated with the toggle switch.**
The text surrounded by a black border represents the stochastic simulation algorithm integrated into the 3D colony agent-based model.

##### 3. Determination of First Passage Time

Due to the varying and noisy distributions of the toggle, we employ a statistical learning method to determine all cell states. We use a Gaussian mixture model for classification of the binary states (Green or Red states) of cells (see **Fig. S5**). In brief, about 40,000 trajectories were generated using random initial conditions, i.e., we chose a set of random data as the initial conditions of the cells, then, evolved the cells for over 10 generations. The final distribution is used to train the model. Afterwards, we can deploy this model to classify the cell states for all trajectories. The first time the cell state changes from green to red one is the first passage time (FPT). To determine the distribution of FPT, we generate 200,000 trajectories and set the simulation time to 100 doubling time, ensuring that all cells transition from the green to the red state.

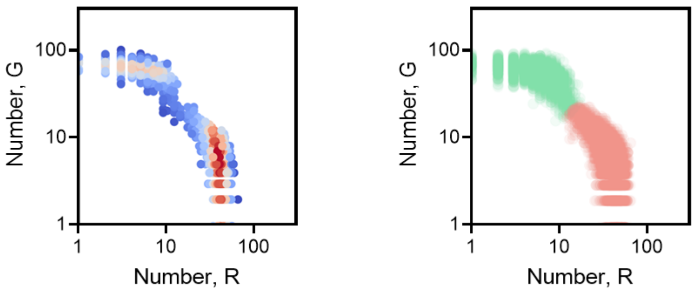

**Fig. S5: Cell state determination**
The left panel displays density scatters of a cell trajectory set, while the right panel shows the binary classified cell states as predicted by a statistical learning model. Red scatters indicate the red state, and green scatters denote the green states.

### Supplementary Figures

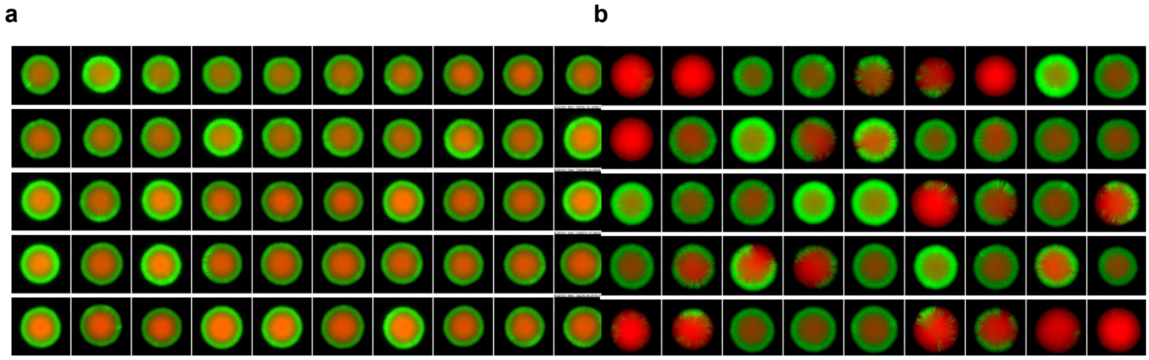

**Fig. S6: Colony patterns from (a) green state single cell and (b) red state single cell.** Forty-five colony patterns grown on RDM glucose agar plates are displayed for each case. Colonies initiated from green state cells exclusively exhibited ring-like patterns, whereas colonies from red state cells displayed diverse patterns.

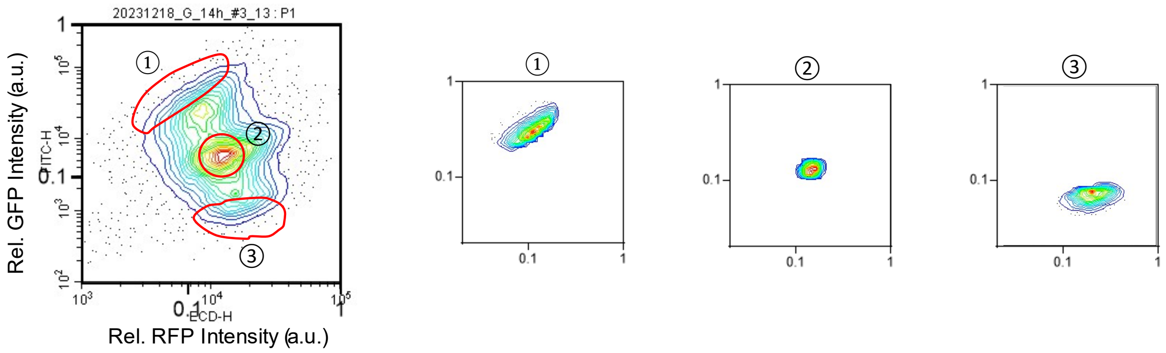

**Fig. S7: FACS gating strategy for bacterial colonies grown on RDM glucose agar.**

Subpopulations ①-③ correspond to the colony periphery, colony interior, and colony top, respectively (see Fig. 2c).

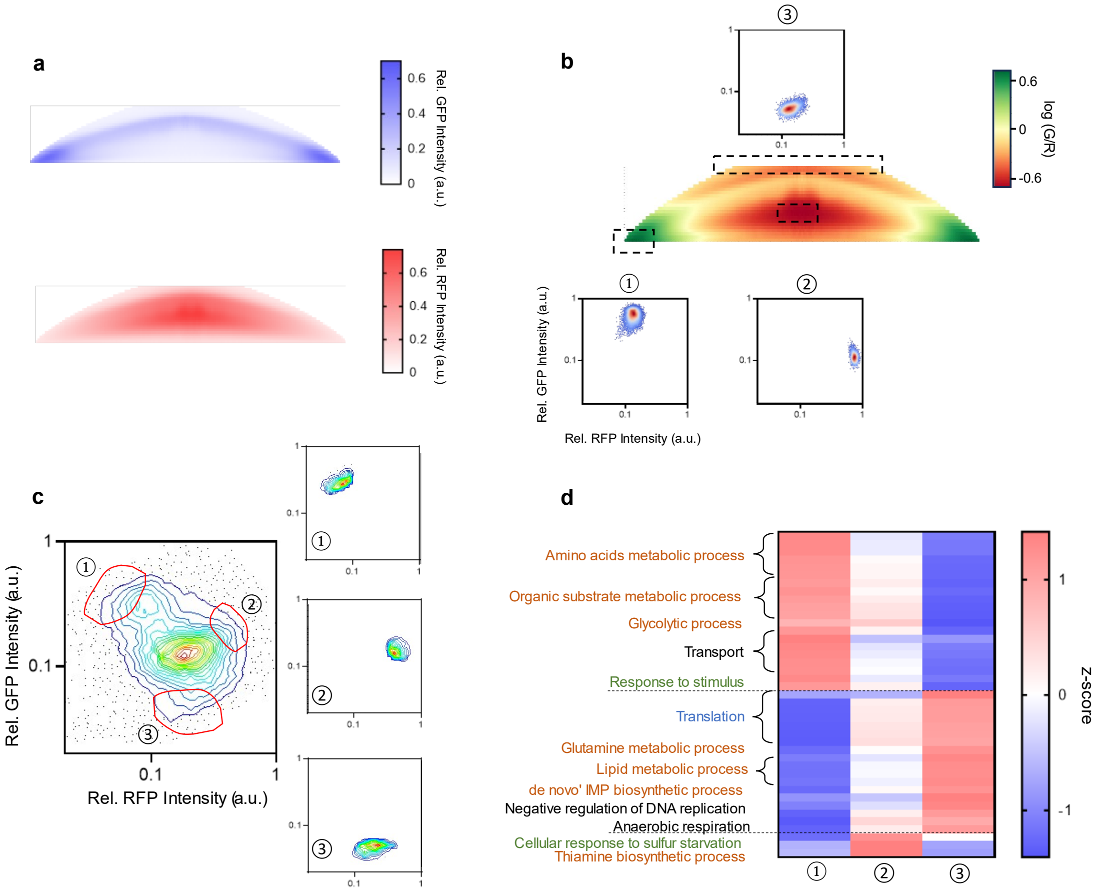

**Fig. S8: Spatial resolved transcriptomes of bacterial colony grown on 1.5% agar supplemented with MOPS glucose medium.**

(a-b) Cross-sectional views depict the spatial distribution of (a) GFP intensity and RFP intensity obtained and reconstructed by two-photon microscopy, and (b) the ratio of GFP intensity to RFP intensity (G/R). Fluorescence intensities were normalized to their corresponding maximal values, and the logarithmic value of the intensity ratio is displayed in panel b for clarity. Green state cells were predominantly observed at the periphery (①), red state cells were located at the top of the colony (③), and co-activated state cells were found at the center (②).

(c) The FACS gating strategy enabled the separation of three distinct subpopulations from different regions of the colony based on their fluorescence intensity profiles.

(d) Metabolic processes, including amino acid synthesis and organic substrate synthesis (see Table S6), were upregulated at the colony periphery. Similar to the case on rich medium (RDM, see Fig. 2c), genes involved in responding to environmental stimuli and membrane transporters were upregulated in this region. Translation-related genes and anaerobic respiration genes were upregulated at the colony top.

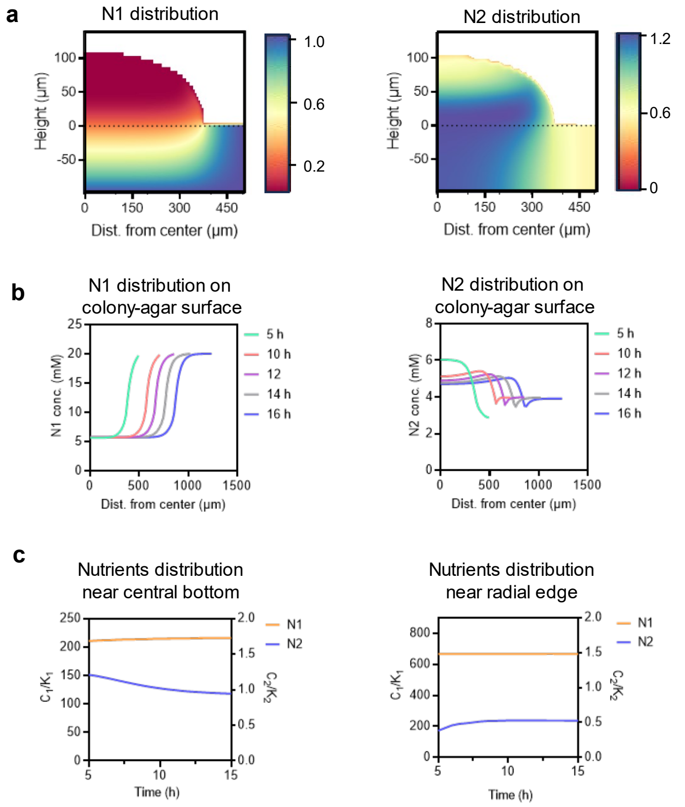

**Fig. S9: Model reveals establishment of stable nutrient distribution at a colony's advancing front.**

(a) Cross-sectional views showing normalized concentrations of nutrients at time t = 5 hours. The left panel indicates the distribution of nutrient N1, and the right panel shows the distribution of nutrient N2.

(b) Nutrient distribution at the colony-agar interface at different times. The left panel represents the distribution of N1, and the right panel indicates the distribution of N2. As growth continues alongside migration patterns, these nutrients reach stable levels.

(c) Dynamics of nutrient concentrations near the central bottom region (left) and radial edges (right) throughout the simulation.

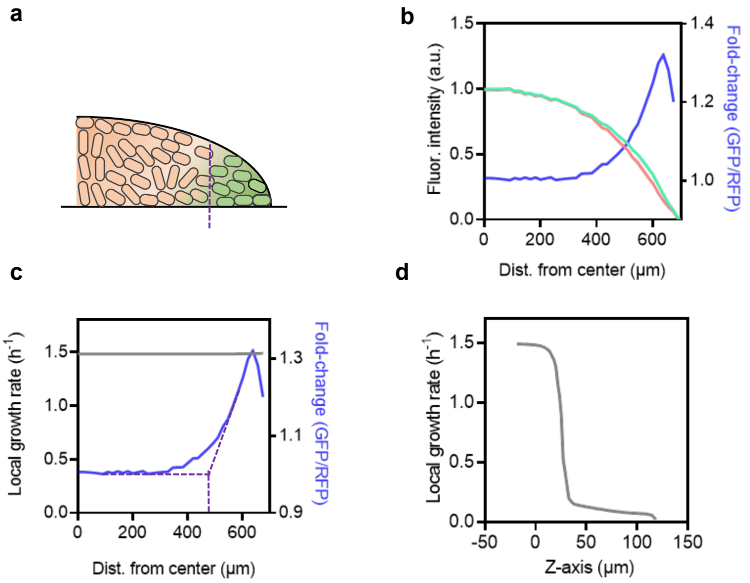

**Fig. S10: Simulation reveals the distribution of fluorescent intensity and local growth rate at an advancing front.**

(a) Schematic of the advancing front of a colony. Cells at the most advanced front of the radial edge remain in the green state, while cells near the central bottom tend to shift toward the red state due to nutrient depletion (monostable state).

(b) Distributions of fluorescent protein (FP) concentrations along the colony radius (green and red lines). Fluorescence intensities are calculated by summing FP concentrations along the z-axis and normalizing by the maximum values. The fold-change (blue dashed line) is calculated as the ratio of GFP to RFP. This exhibits a non-linear trend: as one moves radially from the central bottom (r = 0), the fold-change rapidly increases, reaching a maximum near the colony edge before decreasing. This suggests the dominant GFP expression near the radial edge, resulting in a ring-like pattern when the colony is viewed from the bottom.

(c) Local growth rates along the radius at the colony-agar interface (grey solid line) plotted alongside the fold-change of fluorescence intensities (blue dashed line). Notably, there is no significant variation in local growth rate from the periphery toward the center at the agar surface, indicating that nutrient distribution at the agar surface does not contribute to variations in fluorescence distribution profiles. Extrapolation of the fold-change from its maximum to the value at the central bottom identifies a point of cell fate transition (purple dashed line).

(d) The local growth along the z-axis at the cell fate transition point (purple dashed line in panel a and c) sharply decreases from the bottom to the top of the colony, crossing the bifurcation point of the bistability of the toggle switch.

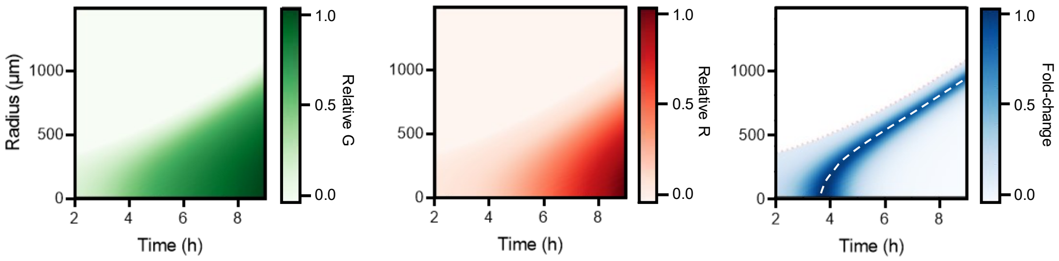

**Fig. S11: Model reveals a constant band width during the establishment phase when colony radius increases linearly with time.**
The radial profiles of relative G state intensity and relative R state intensity at different times are recorded. The right panel illustrates the changes in the fold-change (G/R) over time, characterizing a constant band where the G state expression predominates. The white dashed line indicates the collection of peak values of fold-changes. Time in the figures indicates the simulation time, starting from the condition where the radius is approximately 300 μm.

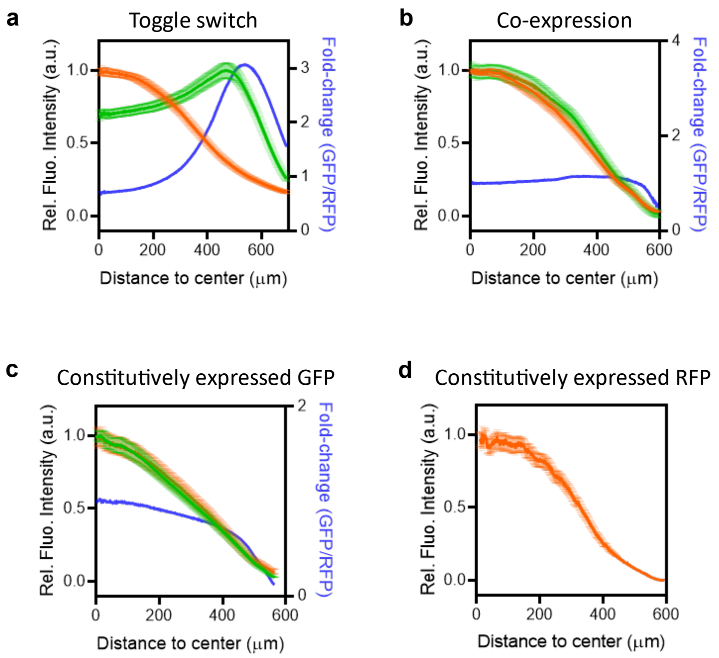

**Fig. S12: Characterization of band width using wide-field microscope images.**

Bacterial colony patterns harboring (a) a synthetic toggle switch (LO1 in Table S2), (b) co-expression (pHC3_mRFP_mVenus in Table S2), (c) constitutively expressed GFP (pECJ3_M5_delta_ptrc in Table S2), and (d) constitutively expressed RFP (pECJ3_M5_delta_PLtetO in Table S2) imaged by wide-field microscopy. Only the pattern programmed by the synthetic toggle switch exhibits a prominent peak when calculating the fold-change (ratio of GFP intensity to RFP intensity, GFP/RFP). Thus, the distance between the colony edge and the GFP/RFP peak is calculated to characterize the ring-like pattern programmed by the toggle switch.

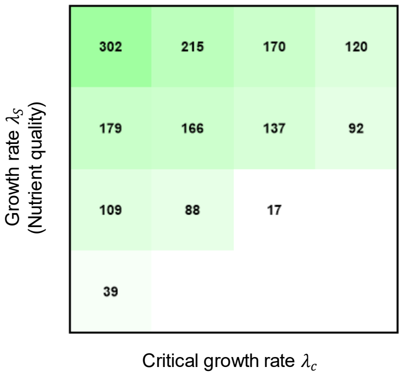

**Fig. S13: The bivariate dependency of ring width on the critical transition growth rate $\lambda_c$ and the maximum growth rate $\lambda_s$.**

Values represent the mean band width in μm characterized under different conditions.

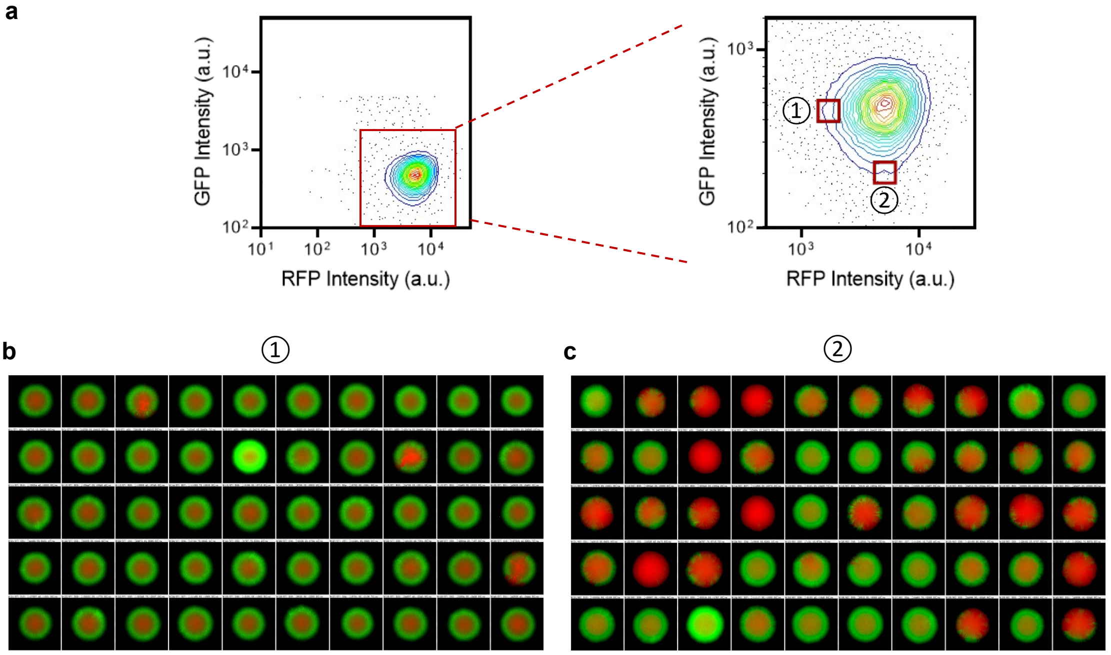

**Fig. S14: Colony patterns formed from initial red state cells within different subpopulations.** Cells were sorted using FACS with “single” sort mode. Fifty colony patterns grown on RDM glucose agar plates are displayed for each case.

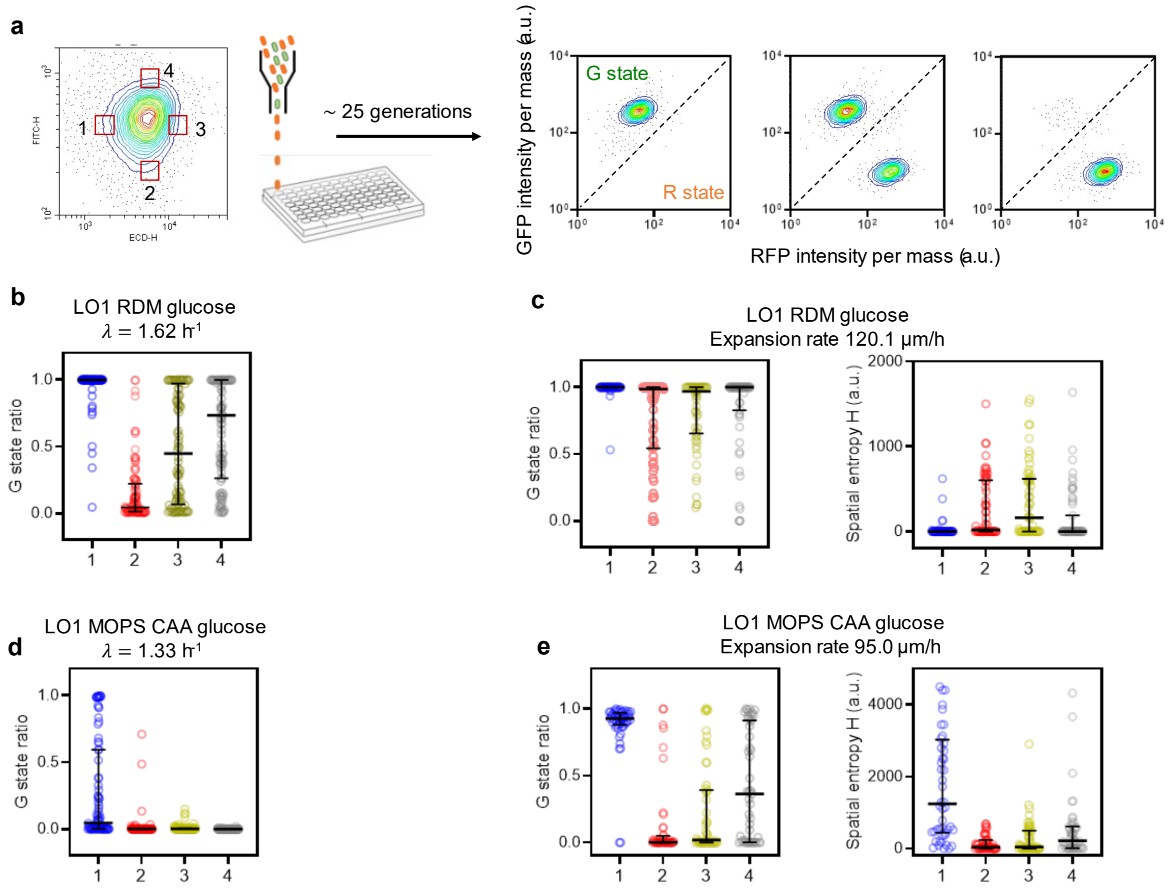

**Fig. S15: Experimental evaluation of the effects of state transition probability and expansion rate in a range expansion system.**

(a) Similar to the description in Extended Data Fig. 7, the difference in state transition probability was monitored by the G state ratio after approximately 25 generations of growth in liquid culture from a single R state cell sorted from different initial conditions of gene expression. The final $OD_{600}$ was 0.2-0.4, indicating cells were still in the exponential growth phase. The G state ratio was analyzed by flow cytometry, and three typical scenarios are displayed in the right panel.

(b) and (d) show the initial condition dependent G state ratio grown in RDM glucose medium and MOPS CAA glucose medium, respectively. (c) and (e) display the G state ratio and spatial entropy of the expansion frontiers of colony patterns in these two media as the growing substate.

This analysis illustrates the significance of both spontaneous state transition probability and range expansion in the formation of pattern symmetry breaking. 90-96 samples were collected for each case in liquid culture and 50-60 samples were analyzed for each of the case of colony pattern. The lines in the plots indicate the median and interquartile ranges.

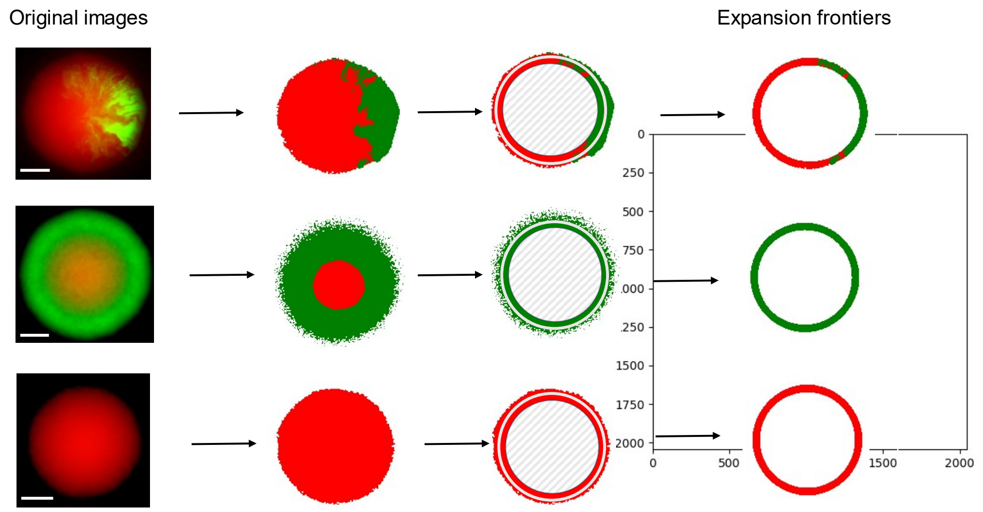

**Fig. S16: Schematic of image processing for statistical analysis.**

Bands with a width of 60 pixels, equivalent to 97.5 μm, positioned 300 to 360 pixels away from the colony center, were utilized for calculating the G state ratio and spatial entropy (see Methods).

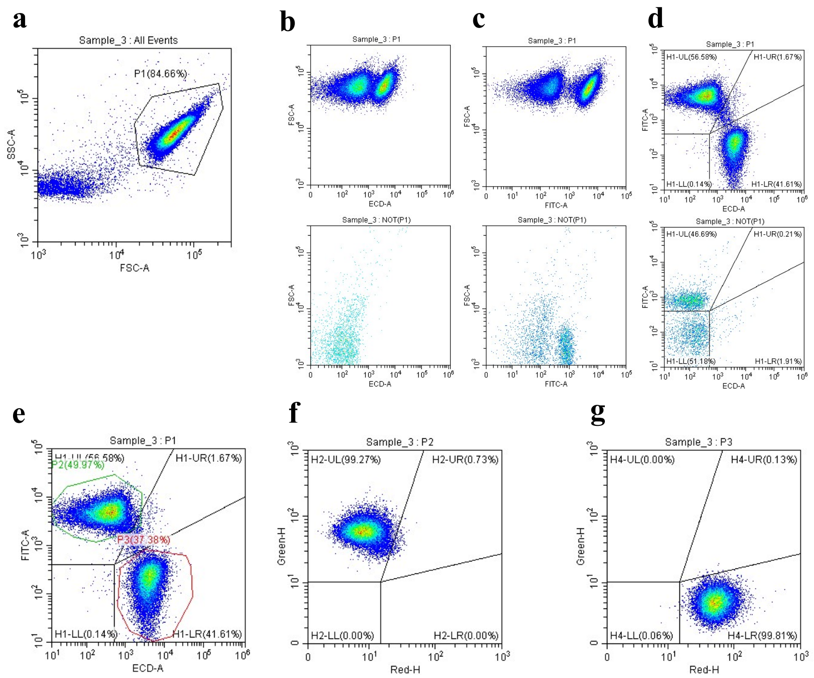

**Fig. S17: Gating strategy used in flow cytometry analysis.**

(a) Particles in P1 were regarded as bacterial cells. Cells harboring the mutual repressive circuit can display a high GFP expression state, a high RFP expression state, or a co-expression state in some cases. An example with a mixture of states is indicated. (b) FSC-A against FITC-A plots for P1 and (c) FSC-A against ECD-A plots for NOT(P1) subpopulations. (d) Plot of FITC-A/ECD-A, divided into subpopulations corresponding to high GFP expression, co-expression, high RFP expression, and non-expression states. (e) Cells in P2 and P3 subpopulations were considered as the green state and the red state, respectively. (f-g) Plots of GFP intensity per mass against RFP intensity per mass using customized parameters (see Methods) show two distinct states.

### Supplementary Tables

**Table S1:**

| **Strain** | **Parent** | **Genotype** | **Source** |
|---|---|---|---|
| NCM3722 | | wild-type *E. coli* K12 strain | Liu Lab |
| NH3 | NCM3722 | NCM3722 Δ*lacIZYA* Δ*fliC* | [10] |

**Table S2: Plasmids constructed in this study.**

| Plasmid | Ori | Antibiotic | Function | Source |
|---|---|---|---|---|
| pECJ3 | ColE1 | Kanamycin | Mutual repression circuit | Gift from Dr. James Collins (Addgene, # 75465) [12] |
| pECJ3_LO1 | ColE1 | Kanamycin | Mutual repression circuit | [10] |
| pECJ3_LO2 | ColE1 | Kanamycin | Mutual repression circuit | [10] |
| pECJ3_LO3 | ColE1 | Kanamycin | Mutual repression circuit | [10] |
| pECJ3_LH1 | ColE1 | Kanamycin | Mutual repression circuit | This study |
| pECJ3_M5_delta_ptrc | ColE1 | Kanamycin | Constitutively expressed LacI-GFP | This study |
| pECJ3_M5_delta_PLtetO | ColE1 | Kanamycin | Constitutively expressed TetR-mCherry | This study |
| pHC3_mRFP_mVenus | p15A | Chloramphenicol | Co-expressed mVenus and mRFP | This study |

**Table S3: Chemical components of defined media.**

| Medium Name | Buffer | Carbon source | Other suppl. | Note |
|---|---|---|---|---|
| RDM glucose | MOPS | 0.4% (w/v) glucose | AUCG + EZ | |
| MOPS CAA glucose | MOPS | 0.4% (w/v) glucose | CAA | |
| MOPS 17AA glucose | MOPS | 0.4% (w/v) glucose | 17AA[a] | |
| MOPS glucose | MOPS | 0.4% (w/v) glucose | / | |
| MOPS xylose | MOPS | 0.4% (w/v) xylose | / | |
| MOPS arabinose | MOPS | 0.4% (w/v) arabinose | / | |
| MOPS fructose | MOPS | 0.4% (w/v) fructose | / | |

[a] 17AA refers to amino acids in EZ, excluding asparagine, aspartic acid, and serine, as indicated in Table S4.

**Table S4: Chemical compositions of supplements for growth media.**

| Supplement name | Compound | Final conc. (mM) |
|---|---|---|
| AUCG | Adenine (Sigma-Aldrich A8626) | 0.2 |
| | Uracil (Sigma-Aldrich U0750) | 0.2 |
| | Cytosine (Sigma-Aldrich C3506) | 0.2 |
| | Guanine (Sigma-Aldrich V900473) | 0.2 |
| EZ | Alanine (Sigma-Aldrich A7469) | 0.8 |
| | Arginine (Sigma-Aldrich V900303) | 5.2 |
| | Asparagine (Sigma-Aldrich V900458) | 0.4 |
| | Aspartic acid (Sigma-Aldrich V900302) | 0.4 |
| | Cysteine (Sigma-Aldrich C6852) | 0.1 |
| | Glutamic acid (Sigma-Aldrich 49601) | 0.6 |
| | Glutamine (Sigma-Aldrich V900419) | 0.6 |
| | Glycine (Sigma-Aldrich G8790) | 0.8 |
| | Histidine (Sigma-Aldrich V900423) | 0.2 |
| | Isoleucine (Sigma-Aldrich V900472) | 0.4 |
| | Leucine (Sigma-Aldrich V900431) | 0.8 |
| | Lysine (Sigma-Aldrich V900438) | 0.4 |
| | Methionine (Sigma-Aldrich V900487) | 0.2 |
| | Phenylalanine (Sigma-Aldrich V900489) | 0.4 |
| | Proline (Sigma-Aldrich V900338) | 0.4 |
| | Serine (Sigma-Aldrich V900406) | 10 |
| | Threonine (Sigma-Aldrich V900466) | 0.4 |
| | Tryptophane (Sigma-Aldrich V900470) | 0.1 |
| | Tyrosine (Sigma-Aldrich V900426) | 0.2 |
| | Valine (Sigma-Aldrich V900465) | 0.6 |
| | Thiamine (Sigma-Aldrich T4625) | 0.01 |
| | Calcium pantothenate (Aladdin C110508) | 0.01 |
| | *p*-aminobenzoic acid (Sigma-Aldrich 100536) | 0.01 |
| | *p*-hydroxybenzoic acid (Sigma-Aldrich V900794) | 0.01 |
| | 2,3-dihydroxybenzoic acid (Sigma-Aldrich 126209) | 0.01 |

#### Table S5: Experimental data from Fig. 2c.

| | | Edge | Inner | Top |
|---|---|---|---|---|
| Metabolic process | Arginine catabolic process to succinate/glutamate | 1.305 | -0.180 | -1.125 |
| | Hypoxanthine metabolic process | 1.049 | 0.296 | -1.346 |
| | 6-sulfoquinovose(1-) catabolic process | 0.894 | 0.502 | -1.396 |
| Response to stimulus | Cellular response to reactive oxygen species | 1.134 | 0.164 | -1.299 |
| | Response to virus | 0.873 | 0.528 | -1.400 |
| | Phage shock | 0.529 | 0.871 | -1.400 |
| Transport | Oligopeptide import across plasma membrane | 0.764 | 0.649 | -1.413 |
| | Phenylalanine transport | 0.296 | 1.049 | -1.346 |
| | Isoleucine biosynthetic process | 0.102 | 1.171 | -1.272 |
| | Tripeptide import across plasma membrane | 0.087 | 1.179 | -1.266 |
| | Heme transmembrane transport | -0.310 | 1.350 | -1.040 |
| | L-glutamate transmembrane transport | -0.439 | 1.384 | -0.945 |
| | L-aspartate import across plasma membrane | -0.445 | 1.385 | -0.940 |
| | Glycine transport | -0.573 | 1.406 | -0.833 |
| | Maltodextrin transmembrane transport | -0.693 | 1.414 | -0.721 |
| Metabolic process | Homocysteine metabolic process | -0.955 | 1.381 | -0.425 |
| | GMP biosynthetic process | -1.057 | -0.285 | 1.342 |
| | Ribonucleoside triphosphate biosynthetic process | -1.069 | -0.267 | 1.336 |
| | Ornithine metabolic process | -1.078 | -0.254 | 1.332 |
| | Arginine biosynthetic process | -1.079 | -0.251 | 1.331 |
| | Glutamine metabolic process | -1.091 | -0.234 | 1.325 |
| | Nucleobase biosynthetic process | -1.093 | -0.231 | 1.324 |
| | Pyrimidine ribonucleotide biosynthetic process | -1.105 | -0.211 | 1.317 |
| | 'de novo' IMP biosynthetic process | -1.111 | -0.203 | 1.313 |
| | Organic phosphonate catabolic process | -1.122 | -0.185 | 1.307 |
| | Regulation of cell shape | -1.089 | -0.237 | 1.326 |
| Translation | Ribosome biogenesis | -1.110 | -0.204 | 1.314 |
| | tRNA aminoacylation | -1.126 | -0.178 | 1.304 |
| | tRNA modification | -1.169 | -0.104 | 1.274 |
| Transport | Organic phosphonate transport | -1.138 | -0.158 | 1.296 |
| | Sulfur compound biosynthetic process | -1.143 | -0.149 | 1.293 |
| | L-arginine import across plasma membrane | -1.155 | -0.129 | 1.284 |
| | Protein localization to membrane | -1.183 | -0.080 | 1.263 |
| | Phospholipid biosynthetic process | -1.244 | 0.039 | 1.205 |

#### Table S6: Experimental data from Fig. S8d.

| | | Edge | Inner | Top |
|---|---|---|---|---|
| Amino acids synthesis and metabolic process | Tryptophan biosynthetic process | 1.321 | -0.224 | -1.097 |
| | Isoleucine biosynthetic process | 1.301 | -0.171 | -1.131 |
| | Valine biosynthetic process | 1.300 | -0.169 | -1.132 |
| | Aspartate metabolic process | 1.198 | 0.053 | -1.250 |
| | Leucine biosynthetic process | 1.171 | 0.102 | -1.273 |
| | Histidine biosynthetic process | 1.166 | 0.110 | -1.276 |
| Organic substrate metabolic process | Dicarboxylic acid biosynthetic process | 1.116 | 0.194 | -1.310 |
| | Glycogen metabolic process | 1.193 | 0.062 | -1.255 |
| | NADP metabolic process | 1.108 | 0.207 | -1.315 |
| | Glutathione metabolic process | 1.046 | 0.301 | -1.347 |
| | Pyridine nucleotide biosynthetic process | 1.024 | 0.333 | -1.357 |
| | NAD metabolic process | 0.829 | 0.578 | -1.407 |
| Glycolytic process | Glycolytic process (ADP/ATP metabolic process) | 1.152 | 0.134 | -1.286 |
| Transport | Intermembrane phospholipid transfer | 1.400 | -0.526 | -0.874 |
| | Molybdat/tungstatee ion transport | 1.325 | -0.234 | -1.091 |
| | Glucose import across plasma membrane | 1.297 | -0.160 | -1.137 |
| | L-histidine import across plasma membrane | 1.255 | -0.062 | -1.193 |
| | Phosphate ion transport | 1.235 | -0.021 | -1.214 |
| Response to stimulus | Response to osmotic stress | 1.306 | -0.182 | -1.123 |
| | Cellular response to oxidative stress | 1.143 | 0.150 | -1.293 |
| Translation | Protein refolding | -0.760 | -0.652 | 1.413 |
| | tRNA modification | -1.307 | 0.185 | 1.122 |
| | Cytoplasmic translation | -1.322 | 0.226 | 1.096 |
| | Ribosomal small subunit biogenesis | -1.325 | 0.234 | 1.091 |
| | Regulation of translation | -1.346 | 0.296 | 1.050 |
| | Ribosome assembly | -1.351 | 0.315 | 1.037 |
| | Ribosomal large subunit assembly | -1.363 | 0.356 | 1.007 |
| Glutamine metabolic process | Glutamine metabolic process | -1.238 | 0.027 | 1.211 |
| Biosynthesis | Pyruvate catabolic process | -1.092 | -0.233 | 1.324 |
| | Phospholipid biosynthetic process | -1.187 | -0.073 | 1.260 |
| | Fatty acid biosynthetic process | -1.197 | -0.053 | 1.250 |
| | Lipooligosaccharide/glycolipid biosynthetic process | -1.157 | -0.126 | 1.283 |
| | de novo' IMP biosynthetic process | -1.252 | 0.056 | 1.196 |
| Transport | Spermidine transmembrane transport | -0.909 | -0.483 | 1.393 |
| Response to stimulus | Response to heat | -1.061 | -0.279 | 1.340 |
| | Response to cold | -1.308 | 0.187 | 1.120 |
| | Negative regulation of DNA replication | -1.384 | 0.440 | 0.944 |
| | Anaerobic respiration | -1.314 | 0.203 | 1.111 |
| | Cellular response to sulfur starvation | -1.230 | 1.220 | 0.010 |
| | Thiamine biosynthetic process | -0.646 | 1.413 | -0.766 |
| | Putrescine catabolic process | -0.589 | 1.408 | -0.819 |

**Table S7: Growth rate**

| Growth medium | Strain | Cell state | Growth rate ($h^{-1}$) |
|---|---|---|---|
| RDM glucose | LH1 | R | 1.594 |
| | | G | 1.553 |
| | LO1 | R | 1.618 |
| | | G | 1.561 |
| | LO2 | R | 1.574 |
| | | G | 1.605 |
| | LO3 | G | 1.563 |
| MOPS 17AA glucose | LH1 | R | 1.330 |
| | | G | 1.304 |
| | LO1 | R | 1.333 |
| | | G | 1.320 |
| | LO2 | R | 1.330 |
| | | G | 1.299 |
| | LO3 | G | 1.300 |
| MOPS glucose | LH1 | R | 0.966 |
| | | G | 0.913 |
| | LO1 | R | 0.909 |
| | | G | 0.878 |
| | LO2 | R | 0.917 |
| | | G | 0.822 |
| | LO3 | G | 0.864 |
| MOPS fructose | LH1 | R | 0.774 |
| | | G | 0.775 |
| | LO1 | R | 0.782 |
| | | G | 0.734 |
| | LO2 | R | 0.772 |
| | | G | 0.681 |
| | LO3 | R | 0.750 |
| | | G | 0.705 |
| MOPS xylose | LH1 | R | 0.736 |
| | | G | 0.725 |
| | LO1 | R | 0.745 |
| | | G | 0.676 |
| | LO2 | R | 0.726 |
| | | G | 0.617 |
| | LO3 | R | 0.681 |
| | | G | 0.645 |

#### Table S8: Functional parts used in this study.

| Part name | Sequence (5'→ 3') | Function |
|---|---|---|
| Ptrc | TTGACAATTAATCATCCGGCTCGTATAATGTGTGGAATTGTGAGC<br>GGATAACAA | Promoter |
| Ptrc_LO1 | TTGACAATTAATCATCCGGCTCGTATAATGTGTGGAATTGTTACTC<br>GCTCACAT | Promoter |
| Ptrc_LO2 | TTGACAATTAATCATCCGGCTCGTATAATGTGTGGAAATGTGAGC<br>GAGTAACAA | Promoter |
| Ptrc_LO3 | TTGACAATTAATCATCCGGCTCGTATAATGTGTGGAAAATTGTGAG<br>CGCTCACAATT | Promoter |
| Ptrc_LH1 | TTGACAATTAATCATCCGGCTCGTATAATGTGTGGAAAATTCTGAG<br>CGCTCACAATT | Promoter |
| *lacI* | ATGGTGAATGTGAAACCAGTAACGCTGTACGATGTCGCAGAGTAT<br>GCCGGTGTCTCTTATCAGACCGTTTCCCGCGTGGTGAACCAGGC<br>CAGCCACGTTTCTGCGAAAACGCGGGAAAAAGTGGAAGCGGCGA<br>TGGCGGAGCTGAATTACATTCCCAACCGCGTGGCACAACAACTG<br>GCGGGCAAACAGTCGTTGCTGATTGGCGTTGCCACCTCCAGTCT<br>GGCCCTGCACGCGCCGTCGCAAATTGTCGCGGCGATTAAATCTC<br>GCGCCGATCAACTGGGTGCCAGCGTGGTGGTGTCGATGGTAGAA<br>CGAAGCGGCGTCGAAGCCTGTAAAGCAGCGGTTCACAATCTTCT<br>CGCGCAGCGCGTCAGTGGGCTGATCATTAACTATCCGCTGGATG<br>ACCAGGATGCCATTGCTGTGGAAGCTGCCTGCACTAATGTTCCG<br>GCGTTATTTCTTGATGTCTCTGACCAGACACCCATCAACAGTATTA<br>TTTTCTCCCATGAAGACGGTACGCGACTGGGCGTGGAGCATCTG<br>GTCGCATTGGGTCACCAGCAAATCGCGCTGTTAGCGGGCCCATT<br>AAGTTCTGTCTCGGCGCGTCTGCGTCTGGCTGGCTGGCATAAAT<br>ATCTCACTCGCAATCAAATTCAGCCGATAGCGGAACGGGAAGGC<br>GACTGGAGTGCCATGTCCGGTTTTCAACAAACCATGCAAATGCTG<br>AATGAGGGCATCGTTCCCACTGCGATGCTGGTTGCCAACGATCA<br>GATGGCGCTGGGCGCAATGCGCGCCATTACCGAGTCCGGGCTG<br>CGCGTTGGTGCGGACATCTCGGTAGTGGGATACGACGATACCGA<br>AGACAGCTCATGTTATATCCCGCCGTTAACCACCATCAAACAGGA<br>TTTTCGCCTGCTGGGGCAAACCAGCGTGGACCGCTTGCTGCAAC<br>TCTCTCAGGGCCAGGCGGTGAAGGGCAATCAACTGTTGCCCGTC<br>TCACTGGTGAAAAGAAAAACCACCCTGGCTCCCAATACGCAAACC<br>GCCTCTCCCCGCGCGTTGGCCGATTCATTAATGCAACTGGCACG<br>ACAGGTTTCCCGACTGGAAAGCGGGCAGGCGGCGAACAAAAACG<br>AAGAAAACACCAACGAAGTGCCGACCTTTATGCTGAACGCGGGC<br>CAGGCGAACAGAAGACGAGTTTAA | CDS |
| *tetR* | ATGTCTCGTTTAGATAAAAGTAAAGTGATTAACAGCGCATTAGAGC<br>TGCTTAATGAGGTCGGAATCGAAGGTTTAACAACCCGTAAACTCG<br>CCCAGAAGCTAGGTGTAGAGCAGCCTACATTGTATTGGCATGTAA<br>AAAATAAGCGGGCTTTGCTCGACGCCTTAGCCATTGAGATGTTAG<br>ATAGGCACCATACTCACTTTTGCCCTTTAGAAGGGGAAAGCTGGC<br>AAGATTTTTTACGTAATAACGCTAAAAGTTTTAGATGTGCTTTACTA<br>AGTCATCGCGATGGAGCAAAAGTACATTTAGGTACACGGCCTACA<br>GAAAAACAGTATGAAACTCTCGAAAATCAATTAGCCTTTTTATGCC<br>AACAAGGTTTTTCACTAGAGAATGCATTATATGCACTCAGCGCTGT<br>GGGGCATTTTACTTTAGGTTGCGTATTGGAAGATCAAGAGCATCA<br>AGTCGCTAAAGAAGAAAGGGAAACACCTACTACTGATAGTATGCC<br>GCCATTATTACGACAAGCTATCGAATTATTTGATCACCAAGGTGCA<br>GAGCCAGCCTTCTTATTCGGCCTTGAATTGATCATCTGCGGATTA<br>GAAAAACAACTTAAATGTGAAAGTGGGTCTTGA | CDS |
| *mCherry* | ATGGTGAGCAAGGGCGAGGAGGATAACATGGCCATCATCAAGGA<br>GTTCATGCGCTTCAAGGTTCACATGGAGGGCTCCGTGAACGGCC<br>ACGAGTTCGAGATCGAGGGCGAGGGCGAGGGCCGCCCCTACGA<br>GGGCACCCAGACCGCCAAGCTGAAGGTGACCAAGGGTGGCCCC<br>CTGCCCTTCGCCTGGGACATCCTGTCCCCTCAGTTCATGTACGG<br>CTCCAAGGCCTACGTGAAGCACCCCGCCGACATCCCCGACTACT<br>TGAAGCTGTCCTTCCCCGAGGGCTTCAAGTGGGAGCGCGTGATG<br>AACTTCGAGGACGGCGGCGTGGTGACCGTGACCCAGGACTCCTC<br>CCTGCAAGACGGCGAGTTCATCTACAAGGTGAAGCTGCGCGGCA<br>CCAACTTCCCCTCCGACGGCCCCGTAATGCAGAAGAAGACTATG<br>GGCTGGGAGGCCTCCTCCGAGCGGATGTACCCCGAGGACGGCG<br>CGCTGAAGGGCGAGATCAAGCAGAGGCTGAAGCTGAAGGACGG<br>CGGCCACTACGACGCTGAGGTCAAGACCACCTACAAGGCCAAGA<br>AGCCCGTGCAACTGCCCGGCGCGTACAACGTCAACATCAAGTTG<br>GACATCACCTCCCACAACGAGGACTACACCATCGTGGAACAGTA<br>CGAACGCGCCGAGGGCCGCCACTCCACCGGCGGCATGGACGAG<br>CTGTATAAGTAA | CDS |
| *kanR* | ATGATTGAACAAGATGGATTGCACGCAGGTTCTCCGGCGGCTTG<br>GGTGGAGAGGCTATTCGGCTATGACTGGGCACAACAGACAATCG<br>GCTGCTCTGATGCCGCCGTGTTCCGGCTGTCAGCGCAGGGTCGC<br>CCGGTTCTTTTTGTCAAGACCGACCTGTCCGGTGCCCTGAATGAA | CDS |

| | | |
|---|---|---|
| | CTGCAAGACGAGGCAGCGCGGCTATCGTGGCTGGCCACGACGG<br>GCGTTCCTTGCGCGGCTGTGCTCGACGTTGTCACTGAAGCGGGA<br>AGGGACTGGCTGCTATTGGGCGAAGTGCCGGGGCAGGATCTCCT<br>GTCATCTCACCTTGCTCCTGCCGAGAAAGTATCCATCATGGCTGA<br>TGCAATGCGGCGGCTGCATACGCTTGATCCGGCTACCTGCCCAT<br>TCGACCACCAAGCGAAACATCGCATCGAGCGAGCACGTACTCGG<br>ATGGAAGCCGGTCTTGTCGATCAGGATGATCTGGACGAAGAGCA<br>TCAGGGGCTCGCGCCAGCCGAACTGTTCGCCAGGCTCAAGGCG<br>CGTATGCCCGACGGCGAGGATCTCGTCGTGACCCACGGCGATG<br>CCTGCTTGCCGAATATCATGGTGGAAAATGGCCGCTTTTCTGGAT<br>TCATCGACTGTGGCCGGCTGGGTGTGGCGGACCGCTATCAGGAC<br>ATAGCGTTGGCTACCCGTGATATTGCTGAAGAGCTTGGCGGCGA<br>ATGGGCTGACCGCTTCCTCGTGCTTTACGGTATCGCCGCTCCCG<br>ATTCGCAGCGCATCGCCTTCTATCGCCTTCTTGACGAGTTCTTCT<br>GA | |
| ColE1 | TTGAGATCCTTTTTTTCTGCGCGTAATCTGCTGCTTGCAAACAAAA<br>AAACCACCGCTACCAGCGGTGGTTTGTTTGCCGGATCAAGAGCT<br>ACCAACTCTTTTTCCGAAGGTAACTGGCTTCAGCAGAGCGCAGAT<br>ACCAAATACTGTCCTTCTAGTGTAGCCGTAGTTAGGCCACCACTT<br>CAAGAACTCTGTAGCACCGCCTACATACCTCGCTCTGCTAATCCT<br>GTTACCAGTGGCTGCTGCCAGTGGCGATAAGTCGTGTCTTACCG<br>GGTTGGACTCAAGACGATAGTTACCGGATAAGGCGCAGCGGTCG<br>GGCTGAACGGGGGGTTCGTGCAGACAGCCCAGCTTGGAGCGAA<br>CGACCTACACCGAACTGAGATACCTACAGCGTGAGCTATGAGAAA<br>GCGCCACGCTTCCCGAAGGGAGAAAGGCGGACAGGTATCCGGT<br>AAGCGGCAGGGTCGGAACAGGAGAGCGCACGAGGGAGCTTCCA<br>GGGGGAAACGCCTGGTATCTTTATAGTCCTGTCGGGTTTCGCCA<br>CCTCTGACTTGAGCGTCGATTTTTGTGATGCTCGTCAGGGGGGC<br>GGAGCCTATGGAAA | CDS |

The characters underscored represent the repressors’ operators.

**Table S9: Parameters used in model.**

| Parameters | Value | Unit | Description | Notes |
|---|---|---|---|---|
| $\delta_t$ | 5.0E-5 | h | Time step to simulate the phase field | This study |
| $N_t$ | 60 | - | Scale for time step to simulate nutrients evolution | This study |
| $\nu_0$ | (10.0, 50.0) | - | Viscosity | This study[a] |
| $D$ | 110 | $\mu m^2/s$ | Diffusion coefficient of nutrients, estimated from other study. | [6] |
| $\epsilon$ | 16 | μm | Width of the interface between air and colony | This study[a] |
| $\Gamma$ | 80.0 | - | Lagrange multiplier for phase field evolution | This study[a] |
| $\gamma$ | 100.0 | - | Coefficient of surface tension | This study[a] |
| $z_s$ | 16 | μm | The location of the agar surface | This study[a] |
| $\delta$ | 16 | μm | Width of substrate | This study[a] |
| $A$ | 10 | - | The adhesion energy | This study[a] |
| $g$ | 1E7 | - | Coefficient of repulsive term of the interface | This study[a] |
| $\lambda_1^{\max}$ | 1.0 | $h^{-1}$ | Growth rates in the saturated $N_1$, | This study[b] |
| $\lambda_2^{\max}$ | 0.4 | $h^{-1}$ | Growth rates in the saturated $N_2$ | This study[b] |
| $K_1$ | 0.5 | μM | Monod constants of $N_1$ | This study, free parameter, we fix it. |
| $K_2$ | 5.0 | μM | Monod constants of $N_2$, this value was estimated from the experimental data of acetate as a sole carbon source. | [13] |
| $f_1$ | 2750.0 | $\mu M \cdot h^{-1}$ | Uptake flux of $N_1$, this value was estimated from the experimental data of glucose as a sole carbon source. | [14] |
| $f_2$ | 3536.1 | $\mu M \cdot h^{-1}$ | Uptake flux of $N_2$, this value was estimated from the experimental data of acetate as a sole carbon source. | [14] |
| $p_2$ | 1222.2 | $\mu M \cdot h^{-1}$ | Excretion flux of $N_2$, this value was estimated from the experimental data of acetate as a sole carbon source. | [14] |
| $C_{1,0}$ | 20.0 | μM | Initial condition of the concentration of $N_1$. Typical value, the concentration of carbon source in growth media. | This study |
| $C_{2,0}$ | 0.0 | μM | Initial condition of the concentration of $N_2$ | This study |
| $\alpha_G(k_g)$ | - | - | LacI expression rates | This study[b] |
| $\alpha_R(k_g)$ | - | - | TetR expression rates | This study[b] |
| $\tau_G, K_G, n_G$ | - | - | Parameters for the hill function | This study[b] |
| $\tau_R, K_R, n_R$ | - | - | Parameters for the hill function | This study[b] |
| $m$ | 192 | - | Initial grid number in r axis | This study[c] |
| $n$ | 64 | - | Initial grid number in z axis | This study[c] |
| $L_r$ | 350 | μm | Initial right boundary of the r axis | This study[c] |
| $L_z^{bottom}$ | 100 | μm | Initial bottom boundary of the z axis | This study[c] |
| $L_z^{top}$ | 100 | μm | Initial top boundary of the z axis | This study[c] |

Notes:

[a]. There parameters are associated with the mechanical properties of the colony, such as the ratio of colony size to colony radius. We tune these parameters to achieve a best fit with the geometric shape of the colony.

[b]. The gene expression rates are fitted from experimental data.

$\alpha_G(k_g) = 1.0 \cdot k_g \cdot (40.6 + \frac{270.28}{1.0+(k_g/0.66)^{8.65}})$; $\alpha_R(k_g) = 1.0 \cdot k_g \cdot (26.83 + \frac{320.21}{1.0+(k_g/0.47)^{5.9}})$.

Parameters for the toggle switch are variables depending on the simulation conditions.

For strain NH3 pECJ3_LO3, we used the parameters: $(\tau_G, K_G, n_G) = (0.1202, 12.0, 4.0)$; $(\tau_R, K_R, n_R) = (0.015, 18.25, 2.0)$ . $\tau_R, K_R, n_R$ were fixed in all simulations. $\tau_G, K_G$, and $k_g$ are varied depending on conditions.

[c].Since the colony expands from a scale of ~10 μm to ~ 1000 μm, we extend the simulation field along with the expansion of the colony. We maintain a constant grid size where $L_r/m$ remains unchanged and expand the simulation field when colony radius exceeds half of the current $L_r$.

##### Table S10: Oligonucleotides used in the library preparation.

| **Primer for reverse transcription** | **Sequence** | |
|---|---|---|
| RT_primer 1 | AGAATACACGACGCTCTTCCGATCTGTGTGAANNNNNN | |
| RT_primer 2 | AGAATACACGACGCTCTTCCGATCTTTGGTGANNNNNN | |
| RT_primer 3 | AGAATACACGACGCTCTTCCGATCTGTCACAANNNNNN | |
| RT_primer 4 | AGAATACACGACGCTCTTCCGATCTGCGATAANNNNNN | |
| RT_primer 5 | AGAATACACGACGCTCTTCCGATCTTACAGCANNNNNN | |
| **P5 primer** | **Sequence** | **Index** |
| i509 | 5´AATGATACGGCGACCACCGAGATCTACAC**TTGCTTGC**ACACTCTTTCC CTACACGACGCTCTTCCGATCT-3´ | TTGCTTGC |
| i510 | 5´AATGATACGGCGACCACCGAGATCTACAC**GAGAGGTT**ACACTCTTTCC CTACACGACGCTCTTCCGATCT-3´ | GAGAGGTT |
| i511 | 5´AATGATACGGCGACCACCGAGATCTACAC**ACCTGGTT**ACACTCTTTCC CTACACGACGCTCTTCCGATCT-3´ | ACCTGGTT |
| i512 | 5´AATGATACGGCGACCACCGAGATCTACAC**AAGCGGAA**ACACTCTTTC CCTACACGACGCTCTTCCGATCT-3´ | AAGCGGAA |
| i513 | 5´AATGATACGGCGACCACCGAGATCTACAC**CGGAACAA**ACACTCTTTCC CTACACGACGCTCTTCCGATCT-3´ | CGGAACAA |
| i514 | 5´AATGATACGGCGACCACCGAGATCTACAC**GGTAAGCT**ACACTCTTTCC CTACACGACGCTCTTCCGATCT-3´ | GGTAAGCT |
| i515 | 5´AATGATACGGCGACCACCGAGATCTACAC**TGTGGCAT**ACACTCTTTCC CTACACGACGCTCTTCCGATCT-3´ | TGTGGCAT |
| i516 | 5´AATGATACGGCGACCACCGAGATCTACAC**ACTACGGA**ACACTCTTTCC CTACACGACGCTCTTCCGATCT-3´ | ACTACGGA |
| **N8 primer** | **Sequence** | **Index** |
| N801 | CAAGCAGAAGACGGCATACGAGAT**TCGCCTTA**GTCTCGTGGGCTCGG | TAAGGCGA |
| N802 | CAAGCAGAAGACGGCATACGAGAT**CTAGTACG**GTCTCGTGGGCTCGG | CGTACTAG |
| N807 | CAAGCAGAAGACGGCATACGAGAT**GTAGAGAG**GTCTCGTGGGCTCGG | CTCTCTAC |
| N808 | CAAGCAGAAGACGGCATACGAGAT**CAGCCTCG**GTCTCGTGGGCTCGG | CGAGGCTG |
| N810 | CAAGCAGAAGACGGCATACGAGAT**TCCTCTAC**GTCTCGTGGGCTCGG | GTAGAGGA |
| N812 | CAAGCAGAAGACGGCATACGAGAT**CCTGAGAT**GTCTCGTGGGCTCGG | ATCTCAGG |
| N813 | CAAGCAGAAGACGGCATACGAGAT**TAGCGAGT**GTCTCGTGGGCTCGG | ACTCGCTA |
| N814 | CAAGCAGAAGACGGCATACGAGAT**GTAGCTCC**GTCTCGTGGGCTCGG | GGAGCTAC |
